# Vector-borne *Trypanosoma brucei* parasites develop in artificial human skin and persist as skin tissue forms

**DOI:** 10.1101/2021.05.13.443986

**Authors:** Christian Reuter, Fabian Imdahl, Laura Hauf, Ehsan Vafadarnejad, Philipp Fey, Tamara Finger, Heike Walles, Antoine-Emmanuel Saliba, Florian Groeber-Becker, Markus Engstler

## Abstract

Transmission of *Trypanosoma brucei* by tsetse flies involves the deposition of the cell cycle-arrested metacyclic life cycle stage into mammalian skin at the site of the fly’s bite. In the skin, the metacyclic parasites reactivate and differentiate into proliferative trypanosomes before colonizing the host’s blood and tissues. We generated an advanced human skin equivalent and used tsetse flies to naturally infect the artificial skin with trypanosomes. We detailed the chronological order of the parasites’ development in the skin, and found a rapid activation of metacyclic trypanosomes and differentiation to proliferative parasites. Single-cell parasite transcriptomics were used to document the biological events during differentiation and host invasion at five different timepoints. After the establishment of a proliferative trypanosome population in the skin, the parasites entered a reversible quiescent state characterized by slow replication and a strongly reduced metabolism. We termed these quiescent trypanosomes skin tissue forms (STF), a parasite population that may play an important role in maintaining the infection over long time periods and in asymptomatic infected individuals.

## Introduction

Understanding the first steps of an infection is crucial for early intervention to prevent disease establishment and its progression. In a number of vector-borne parasitic diseases, infection is initiated when pre-adapted parasites are injected by the vector into the dermis of the mammalian skin (Killick-Kendrick, 1990, Tyler and Engman, 2001, Menard et al., 2013).

Unicellular parasites of the *Trypanosoma brucei* (*T. brucei*) group cause human and animal African trypanosomiases and are transmitted to the vertebrate host by the tsetse fly (Vickerman et al., 1988). Infection of vertebrates begins when the fly takes a blood-meal, thereby depositing the infectious metacyclic form (MCF) of the parasite into the dermal skin layer. The MCFs are non-dividing and arrested in G1/G0 (Shapiro et al., 1984). Within the dermis, the MCFs are activated by as yet unknown factors, re-enter the cell cycle, and differentiate into proliferative trypanosomes, which morphologically resemble the proliferative mammalian life cycle stage known as the bloodstream form (BSF) (Dwinger et al., 1988). However, the timing and the mechanisms controlling differentiation remain elusive, as does the characterization of the proliferative skin-residing trypanosomes. From skin tissue, parasites migrate via afferent lymphatic vessels to the local draining lymph nodes before they appear in the bloodstream and systemically infect the host (Barry and Emery, 1984, Alfituri et al., 2020).

One of the salient characteristics of the extracellular parasite is a complex immune evasion mechanism based on antigenic variation of its variant surface glycoprotein (VSG) coat (Cross, 1975, Barry and McCulloch, 2001). The only life cycle stage in the fly vector that expresses VSGs on the surface is the metacyclic stage, which is considered beneficial for the establishment of an infection in the mammalian host. While MCFs express metacyclic VSGs, these are replaced during infection by VSG isoforms characteristic of the bloodstream form (Barry et al., 1998).

Recently, a subpopulation of the injected trypanosomes was found to reside and proliferate in the skin at the bite site (Caljon et al., 2016). In addition, the skin is now recognized as a reservoir tissue during disease, as trypanosomes have frequently been found in the skin of infected animals and humans, even in aparasitemic (i.e. no trypanosomes detectable in blood) and asymptomatic individuals (Capewell et al., 2016, Casas-Sanchez and Acosta-Serrano, 2016, Kruger et al., 2018, Capewell et al., 2019, Camara et al., 2020). The skin-dwelling parasites of laboratory mice can be transmitted to tsetse flies (Capewell et al., 2016, Caljon et al., 2016), strongly suggesting a contribution to disease transmission.

A key goal in research on infectious diseases is the ability to recapitulate the virulence-determining processes in suitable model systems. Human skin equivalents have been successfully used as model systems in infection research, e.g. with *Candida albicans* (Kuhbacher et al., 2017), *Staphylococcus aureus* (Reddersen et al., 2019), and helminths (Jannasch et al., 2015). However, a major problem with collagen-based skin equivalents is their susceptibility to fibroblast-mediated contraction during culture. The poor mechanical stability of the dermal component (Feng et al., 2003) makes standardization difficult.

Here, we have developed an advanced, highly standardized skin infection model, which recapitulates key anatomical, cellular, and functional aspects of native human skin. Using tsetse flies, we have successfully recapitulated the natural vector transmission of *T. brucei* parasites and observed a rapid activation of the cell cycle-arrested MCFs upon arrival in the skin. Furthermore, we have detailed their differentiation to proliferative trypanosomes using a combination of flow cytometry, microscopy, and single-cell RNA sequencing (scRNAseq). Unexpectedly, we found that tsetse-transmitted trypanosomes enter, after an initial proliferative phase, a reversible quiescence program in the skin. This skin-residing trypanosome population, here termed skin tissue forms (STFs), is characterized by slow replication, a strongly reduced metabolism, and their metabolic activity and transcriptome differ from actively proliferating trypanosomes in the skin. By mimicking the migration from the skin to the bloodstream, the quiescent phenotype could be reversed and the parasites returned to an active state. The presence of quiescent trypanosomes hiding in the skin could play a potential role in the development of longstanding infections with little to no symptoms.

## Results

### Improved mechanical properties reduce dermal contraction and weight loss in primary human skin equivalents

In this study, an advanced primary human skin equivalent (hereafter referred to as a high-density skin equivalent) was developed and validated as a model for vector-borne trypanosome skin infection (Figure 1 A). For this, we refined a skin equivalent (Reuter et al., 2017) by improving the mechanical stability of the dermal component to reach a high level of standardization and reproducibility. A computer-assisted compression system consisting of a custom-made compression reactor (Figure S1 A and S1 B) and a linear motor (Figure S1 C) was used to compress the dermal component by a factor of 7. The high-density dermal equivalents (hdDEs) were characterized by a diameter of 12 mm, a height of 1 mm, and a calculated collagen concentration of 46.9 mg/ml.

**Figure 1.**
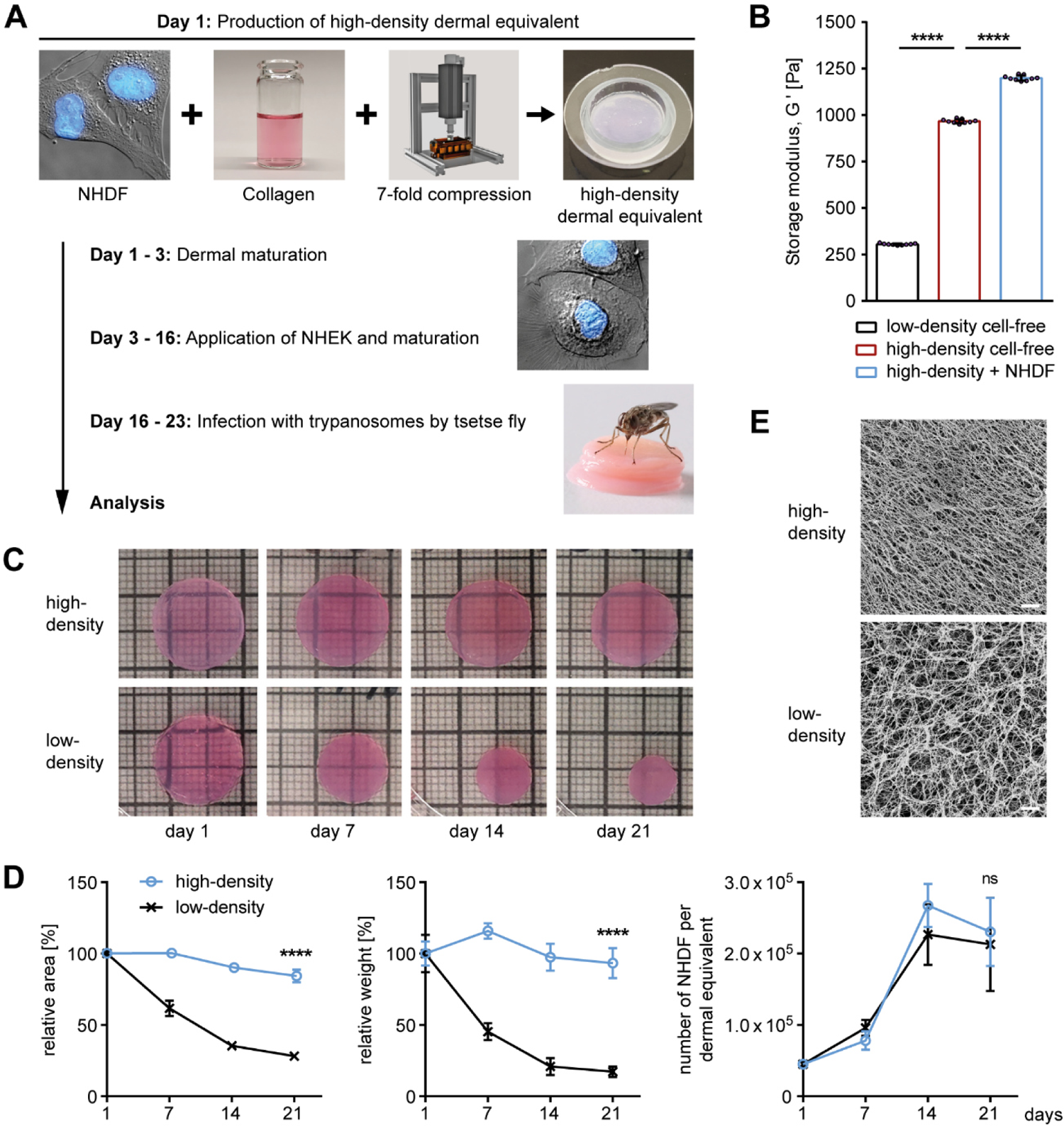
High-density dermal equivalents have improved mechanical properties. **(A)** Normal human dermal fibroblasts (NHDF) were mixed with reconstituted collagen and compressed 7-fold to generate high-density dermal equivalents (hdDEs). Normal human epidermal keratinocytes (NHEK) were added, and the mature high-density skin equivalents were infected with trypanosomes by tsetse flies. **(B)** The storage moduli G′ of NHDF-populated and cell-free, high- and low-density dermal equivalents (ldDEs) indicate improved mechanical properties of hdDEs. The mean ± SD (n = 3 - 6) of G′ was calculated from values in the linear viscoelastic region (Fig. S1 D, gray box). Unpaired t-test, **** p < 0.0001. **(C)** Images of NHDF-populated hdDEs and ldDEs at various culture times show massive shrinkage of ldDEs (day 21: Ø = 12 mm versus 7 mm). **(D)** Quantification of the relative area and weight, and the number of NHDF per dermal equivalent of hdDEs and ldDEs. Graphs represent mean ± SD (n = 9 - 24). Unpaired t-test, **** p < 0.0001, ns, not significant. **(E)** Scanning electron microscopy of the fibrillar collagen ultrastructure of hdDEs and ldDEs show a denser collagen network in hdDEs. Scale bar, 2 µm.

The increased collagen concentration and reduced water content improved the mechanical properties of the hdDEs, as shown by rheology. Strain sweep measurements confirmed a 3-fold higher storage modulus (G′) in the linear viscoelastic region of cell-free hdDEs compared to cell-free, non-compressed low-density dermal equivalents (ldDEs; c_collagen_ = 6.7 mg/ml), indicating stronger viscoelastic strength of the cell-free hdDEs (Figure 1 B and S1 D). The viscoelastic strength was even more pronounced in hdDEs colonized with normal human dermal fibroblasts (NHDF) for 23 days. Furthermore, ldDEs showed an earlier flow point (G′ = G′′) than hdDEs, correlating with a lower structural resistance against shear forces (Figure S1 D).

Long-term comparison of hdDEs and ldDEs confirmed that hdDEs show significantly less contraction and weight loss over time, while having a similar number of proliferative NHDF (Figure 1 C, 1 D, and S1 E). While hdDEs lost only 6.7 % of weight and 15.7 % of area during 3 weeks of culture, ldDEs lost more than 82.7 % and 71.8 %, respectively. Scanning electron microscopy confirmed proper assembly of collagen fibers in both DEs (Figure 1 E). However, in contrast to ldDEs, the collagen fibers in hdDEs showed a denser and more uniform distribution.

### High-density skin equivalents recapitulate key anatomical, cellular, and functional aspects of native human skin

Different culture media were tested to find optimal conditions for co-cultivation of parasites and skin equivalents (Figure S2 A and S2 B). Moreover, to test whether primary cells isolated from different human donors are compatible with each other, high-density skin equivalents (hdSEs) consisting of NHDFs and normal human epidermal keratinocytes (NHEKs) from three different donors were evaluated based on the thickness of the cellular epidermis (Figure S2 B and S2 C). Overall, these results indicate that trypanosomes and hdSEs can be co-cultured with a mixture of trypanosome and skin medium without detrimental effects.

High-density dermal equivalents were colonized with NHEK on day 3 and serial histological analysis showed that the morphology of hdSEs on day 15 resembled that of native human skin, with a fully differentiated epidermis in which the 4 characteristic layers had formed (Figure 2 A, a-d). The epidermis was stable until day 23, and a prolonged culture was possible but ultimately led to a *stratum corneum* with aspects of hyperkeratosis. Thus, in a time window of minimally 9 days, the tissue architecture of the hdSE resembled native human skin.

**Figure 2.**
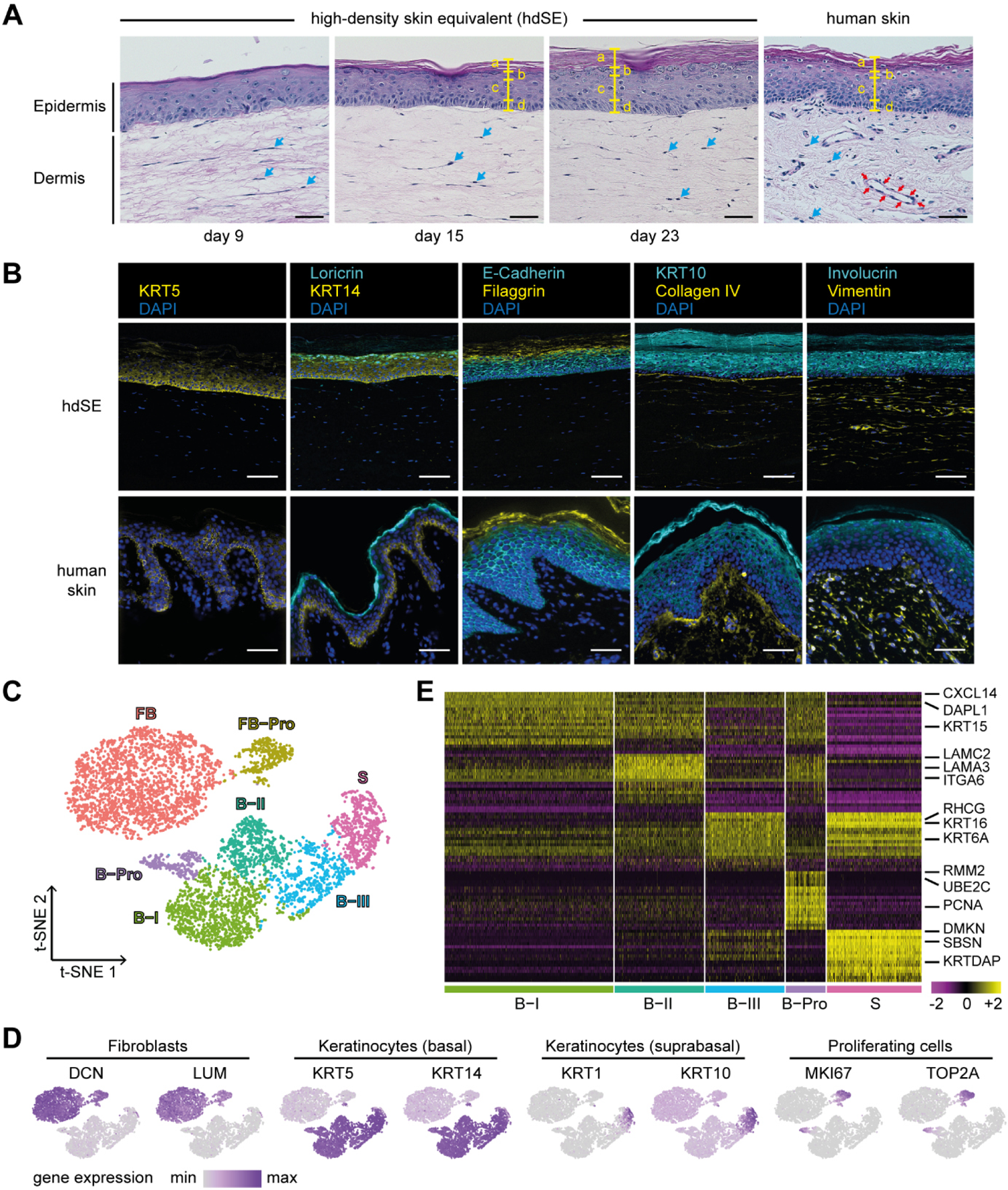
High-density skin equivalents resemble native human skin. **(A)** Hematoxylin and eosin-stained cross sections of high-density skin equivalents (hdSEs) at different culture times in comparison to native human skin. The yellow markings indicate the individual layers of the epidermis: a, *stratum corneum*; b, *stratum granulosum*; c, *stratum spinosum*; d, *stratum basale*. Blue arrows, NHDF. Red arrows, vascular structure. Scale bar, 40 µm. **(B)** Distinctive physiological skin markers were detectable in hdSEs. At day 23, tissues were stained with antibodies against loricrin, E-cadherin, KRT10, involucrin (all cyan), and KRT5, KRT14, filaggrin, collagen IV, and vimentin (all yellow). DNA was visualized with DAPI (blue). Images were acquired by fluorescence microscopy of cross sections of hdSEs. Scale bar, 75 µm. **(C)** t-Distributed stochastic neighbor embedding (t-SNE) plot of 5958 cell transcriptomes derived from two hdSEs at day 23. The major populations of cell types were fibroblasts (FB), proliferating fibroblasts (FB-Pro), basal (B-I - III), suprabasal (S), and proliferating keratinocytes (B-Pro). **(D)** The expression of fibroblast and keratinocyte markers from basal, suprabasal, and proliferating cells. Normalized gene expression levels for each cell were color-coded from gray to purple and overlaid onto the t-SNE plot. **(E)** Heatmap showing the scaled expression levels of the 20 most differentially-expressed genes in each cluster of epidermal keratinocytes. The color key from pink to yellow indicates low to high gene expression levels. Each column represents a single cell, and each row represents an individual gene. Cell-type-specific representative genes are listed to the right.

In addition, a comparative analysis showed that physiological skin markers were expressed in the hdSE at day 23 (Figure 2 B). The presence of epidermal differentiation markers such as KRT5 and KRT14 in the basal layer, KRT10 in the suprabasal layers, and loricrin, filaggrin, and involucrin in the *stratum granulosum* and *stratum corneum* verified an anatomically correct differentiation of the epidermis. The presence of collagen IV demonstrated the presence of basement membrane proteins at the structurally and biochemically complex dermal-epidermal junction. Moreover, the homogeneous distribution of vimentin-positive cells in the dermis indicated the presence of NHDF.

To define the cellular heterogeneity of the hdSE in more detail scRNAseq was performed using the droplet-based 10x Genomics platform (Figure S3 A, (Macosko et al., 2015)). Data analysis of 5958 cells that passed quality control (Figure S3 B and Table S1) resulted in a t-distributed stochastic neighbor embedding (t-SNE) plot displaying 7 clusters with distinct expression profiles (Figure 2 C and Table S1).

All clusters were confirmed by the expression of known marker genes (Figure 2 D). Fibroblasts were identified in two clusters (clusters FB and FB-Pro) by their archetypal markers DCN and LUM (Philippeos et al., 2018). Keratinocytes were detected in 5 clusters and their diversity was mainly due to their degree of differentiation. Basal keratinocytes (clusters B-I, B-II, B-III, and B-Pro) highly expressed KRT5 and KRT14, whereas suprabasal keratinocytes (S) specifically expressed KRT1 and KRT10. Proliferating cells were present in both cell populations. They were identified by their expression of mitotic markers such as MKI67 and TOP2A (clusters FB-Pro and B-Pro).

Differential gene expression analysis of the 5 keratinocyte subclusters revealed 3 clusters of basal keratinocytes (B-I, B-II, and B-III), which is in good agreement with native human skin (Wang et al., 2020). The B-I cluster was, besides KRT5 and KRT14, also characterized by KRT15, DAPL1, and CXCL14. CXCL14 is constitutively expressed in skin and produced by keratinocytes (Meuter and Moser, 2008). Cluster B-II contained mainly genes that are important for the formation of hemidesmosomes and the attachment of the epidermis to the dermis via the basement membrane such as LAMA3, LAMB3, LAMC2, ITGA6, ITGB4, DST, PLEC, COL7A1, and COL17A1 (Figure 2 E, S3 C, and Table S1). Cluster B-III was defined by RHCG, KRT6A, and KRT16, similar to what was found earlier, and it was assumed that these cells could directly differentiate into suprabasal cells (Wang et al., 2020). Suprabasal cells in cluster S expressed known markers for epidermal differentiation such as SBSN or KRTDAP. Cluster B-Pro was enriched for the cell cycle markers UBE2C and PCNA. Furthermore, a detailed analysis of extracellular matrix (ECM)-associated genes revealed a remarkable repertoire of expressed ECM genes in the hdSE (Figure S3 C). Many components of the ECM, including the collagens COL1A1, COL1A2, COL3A1, COL12A1, as well as elastin (ELN), fibronectin (FN1), and fibrillin (FBN1) were expressed in the fibroblast cluster. The same was true for factors involved in matrix assembly, such as MFAP4, SFRP2, LOX, FAP, and ANXA2, as well as for proteins involved in matrix remodeling, such as matrix metallopeptidases (MMP2, MMP14) and TIMP metallopeptidase inhibitors (TIMP1, TIMP3).

### Tsetse-transmitted *T. brucei* parasites can infect the skin equivalents

In order to simulate the infection process in the most natural way, tsetse flies with a mature salivary gland infection were employed to transmit the infective metacyclic form (MCF) to the hdSEs (Figure 3 A and Movie 1). The advantages of vector-borne infection are the natural deposition of the parasites in the skin, and the presence of potentially infection-modulating factors from fly saliva. The tsetse flies penetrated the skin with their proboscis several times, and each incision caused an epidermal wound (Figure S4 A) with a diameter similar to that of the fly’s proboscis (Gibson et al., 2017). The analysis of the skin lesion revealed a complex deposition of the parasites within an intricate bite path network in the dermis (Figure 3 B). On average, one fly injected approximately 4000 infective MCFs into the hdSE (Figure S4 B). Half of the parasites were deposited at a depth between 1 - 2 mm, and the other half almost equally at a depth of 0 - 1 mm and 2 - 3 mm (Figure 3 C).

**Figure 3.**
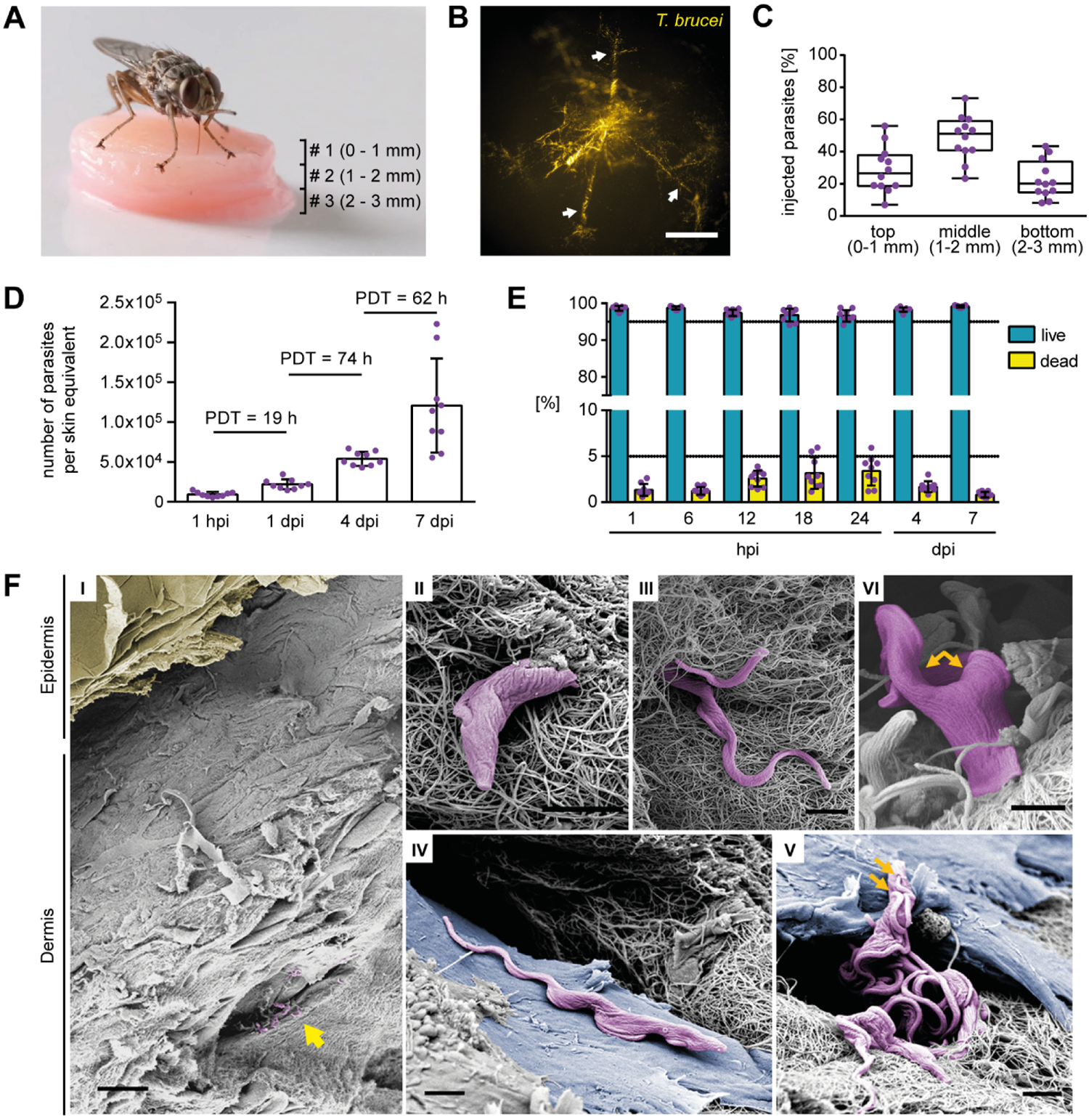
Tsetse-transmitted *T. brucei* parasites establish replicating populations in the skin equivalents. **(A)** Natural infection of skin equivalents with trypanosomes transmitted by a tsetse fly. Three hdSEs were stacked on top of each other (# 1, 2, 3), and tsetse flies with a mature salivary gland infection were subsequently allowed to bite into the stack. **(B)** Stereo-fluorescence microscopy detected tsetse-transmitted metacyclic trypanosomes in the dermis of a skin equivalent immediately after infection. Parasites (yellow) were found in multiple finger-like lesions (white arrows), probably corresponding to the bite path of the tsetse fly. Scale bar, 500 µm. **(C)** Analysis of the injection depth of tsetse-transmitted trypanosomes. Three dermal equivalents with a standardized height of 1 mm were stacked and infected with trypanosomes by tsetse flies. The numbers of parasites in each equivalent were quantified after 1 h and expressed as fractions of the total. Results are shown as median ± IQR (n = 9). **(D)** Cell numbers and population doubling times (PDT) of tsetse-transmitted trypanosomes in skin equivalents at various hours and days post-infection (hpi, dpi) over a 7-day timecourse. Data represent mean ± SD (n = 9). **(E)** Flow cytometry assessment of parasite viability by Calcein-AM staining of infected hdSEs at various timepoints post-infection. Results are shown as mean ± SD (n = 9). **(F)** Scanning electron microscopy of hdSEs 4 days post-infection. (I) Overview showing an intact epidermal layer (colored yellow) and the presence of trypanosomes (colored purple) in the connective tissue of the dermis (yellow arrow). Scale bar, 30 µm. (II + III) Entanglement of trypanosomes with collagen fibers. (IV + V) Parasites were found in close contact with dermal fibroblasts (colored blue). (V + VI) Proliferation was evidenced by the double flagella of trypanosomes (orange arrows). Scale bar, 2 µm (II, III, IV, V, VI).

Within the observation period of 7 days post-infection (dpi), the tsetse-transmitted MCFs increased their total numbers by around two orders of magnitude in the dermis of the hdSEs (Figure 3 D). Measurement of the mean parasite population density (cells/ml) in the hdSEs showed that it increased from an initial 8.24 x 10^4^ cells/ml to 1.07 x 10^6^ cells/ml on day 7. Cell viability assays using Calcein-AM staining (Figure 3 E) detected only low numbers (< 5 %) of dead trypanosomes in the dermis in the hours (hpi) and days (dpi) following infection.

Scanning electron microscopy revealed a high degree of entanglement of skin-dwelling parasites with collagen fibers (Figure 3 F, I - III), and the trypanosomes were frequently found in close contact with dermal fibroblasts (Figure 3 F, IV - V). The presence of dividing trypanosomes was indicated by the observation of cells with a second flagellum (Figure 3 F, V - VI, orange arrows).

### Metacyclic parasites rapidly differentiate into proliferative trypanosomes in the skin equivalents

To measure the timing of the transition of the injected cell cycle-arrested MCFs to proliferative trypanosomes, cell cycle analysis was performed at various timepoints post-infection (Figure 4 A, S4 C, and S4 D). At 12 hpi a 3.6-fold increase of parasites in the S and G2 phases (15.0 %) compared to 6 hpi (4.2 %) was observed (Figure 4 A). The number of dividing parasites further increased after 18 (19.7 %) and 24 hpi (27.1 %). Even after 4 and 7 dpi, the skin-residing parasites remained proliferative (25.9 % and 24.6 %). When analyzed using flow cytometry, the mean fluorescence intensity of the forward scatter width (FSC-W, a measure of cell size) of the injected parasites resembled that of bloodstream form (BSF) culture cells within 24 hpi (Figure 4 B). Microscopy analysis at the same timepoints revealed the presence of cells with a BSF-like morphology (Figure 4 C). Tracking of single parasites in the skin using high-speed microscopy revealed a significant increase in the mean and maximum swimming speeds within the first 24 hpi (Figure 4 D and Movie 2). Moreover, trypanosomes were found in the skin that expressed the VSG AnTat 1.1 isoform, which was on the surface of the parasites used for infection of the tsetse flies (Figure 4 E). However, this was very rare (< 0.1 %) and the earliest time we could detect this VSG was on day 4 post-infection.

**Figure 4.**
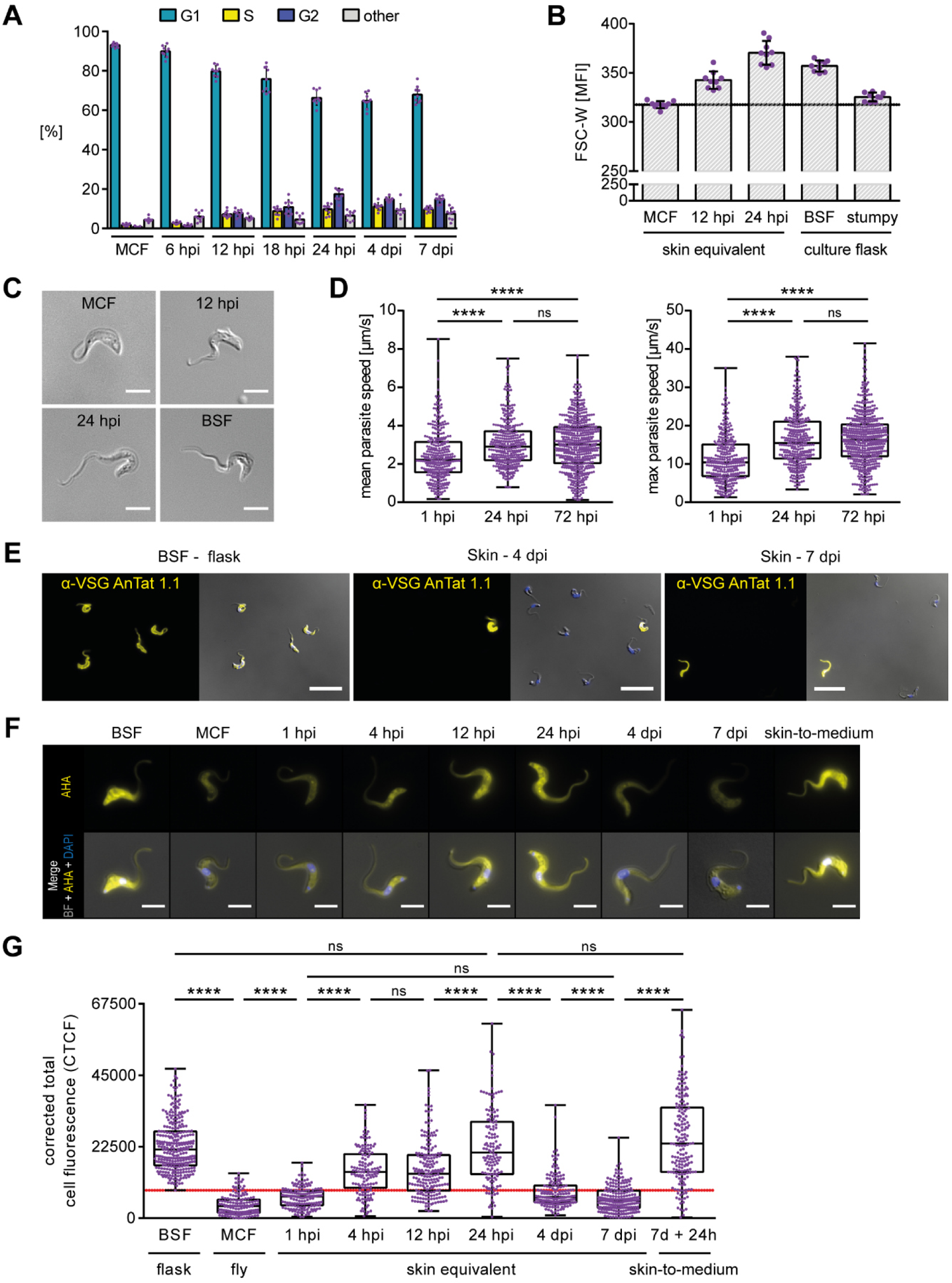
Rapid initiation of proliferative growth by tsetse-transmitted trypanosomes post-infection. **(A)** Flow cytometry quantification of the distribution of cell cycle states using NuclearGreen staining. Isolated metacyclic forms (MCF) and skin-resident trypanosomes at various timepoints post-infection were assayed. Results are shown as mean ± SD (n = 9). **(B)** The mean fluorescence intensity (MFI) of the forward scatter width (FSC-W) was analyzed for MCFs and skin-resident parasites at 12 and 24 hours post-infection (hpi), and compared to bloodstream forms (BSF) and stumpy forms. MCFs were smaller than stumpy forms and both were smaller than BSFs. Results are shown as mean ± SD (n = 9). **(C)** Morphology of an MCF trypanosome and parasites isolated from skin equivalents at 12 and 24 hpi compared to a BSF cell. Scale bar, 5 µm. **(D)** Quantification of the mean (left) and maximum (right) swimming speeds of individual skin-resident trypanosomes at 1, 24, and 72 hpi. Results are shown as median ± IQR (n = 319 - 522). Mann-Whitney U-test, **** p < 0.0001, ns, not significant. **(E)** Immunofluorescence microscopy of skin-derived trypanosomes at 4 and 7 dpi, stained with an antibody against the BSF-specific VSG AnTat 1.1 (yellow channel) and compared to BSFs cultured in flasks. The blue (nuclei, DAPI) and yellow (VSG) channels are shown as overlays with phase contrast image. Scale bar, 20 µm. **(F)** Fluorescence microscopy of trypanosomes isolated at the times indicated post-infection from skin equivalents in comparison to BSFs, MCFs, and skin-parasites at 7 dpi, which were transferred to culture medium for 24 h (skin-to-medium). The fluorescence signal intensity of L-azidohomoalanine (AHA) incorporated into nascent proteins (yellow channel) was measured with the ImageJ/Fiji software to assess protein synthesis rate on the single-parasite level. Cells were counterstained with DAPI (nuclei, blue channel) and both channels were merged with the phase contrast image in the overlays. Scale bar, 5 µm. **(G)** Quantification of protein synthesis rates of BSFs, MCFs, and skin-resident parasites at various timepoints post-infection. Skin-resident parasites were additionally transferred to culture medium for 24 h after 7 dpi (skin-to-medium). The fluorescence intensity of incorporated AHA in individual parasites was measured with ImageJ/Fiji and the corrected total cell fluorescence (CTCF) was calculated. Data are median ± IQR (n = 3; 136 - 292 parasites were analyzed per timepoint). Mann-Whitney U-test, **** p < 0.0001, ns, not significant. The red line defines the threshold for classification as “parasites with low protein synthesis”.

To determine the onset of protein synthesis as indication of MCF activation, the incorporation of a methionine analogue into nascent proteins was quantified (Figure 4 F and 4 G). While MCFs obtained from tsetse flies showed a low protein synthesis rate, a strong 5.4-fold increase was measured in skin trypanosomes within 24 hpi. At that timepoint the rate is comparable to *in vitro* cultured BSFs. The increase in protein synthesis was not linear but stepwise, with a first step between 1 and 4 hpi (2.1-fold), and a second between 12 and 24 hpi (1.5-fold). Already after 1 hpi, the parasites exhibited a 1.8-fold increase in the protein synthesis rate compared to the MCFs that were obtained from flies.

So far, the results strongly suggest that the tsetse-borne cell cycle-arrested MCFs are activated in the hdSE and rapidly re-enter the cell cycle and acquire the size, morphology, motility, and protein synthesis rate characteristic of BSFs.

### Single-parasite RNAseq reveals the programmed activation of metacyclic forms in the skin equivalents

An established developmental hallmark during the differentiation of MCFs is the replacement of the metacyclic VSG coat with VSG isoforms of the BSF. In order to define other programmed changes in gene expression that occur during the developmental transition, the transcriptomes of individual trypanosomes was dissected using Smart-seq2 approach (Picelli et al., 2013). The scRNAseq profiles of MCFs collected from flies with a mature salivary gland infection were compared to those of trypanosomes isolated from hdSEs at 4 different timepoints post-infection (4 hpi, 12 hpi, 24 hpi, and 7 dpi; Figure 5 A). Overall, the transcriptomes of 153 individual parasites were profiled of which 142 parasites passed quality control (Figure S5 A and S5 B).

**Figure 5.**
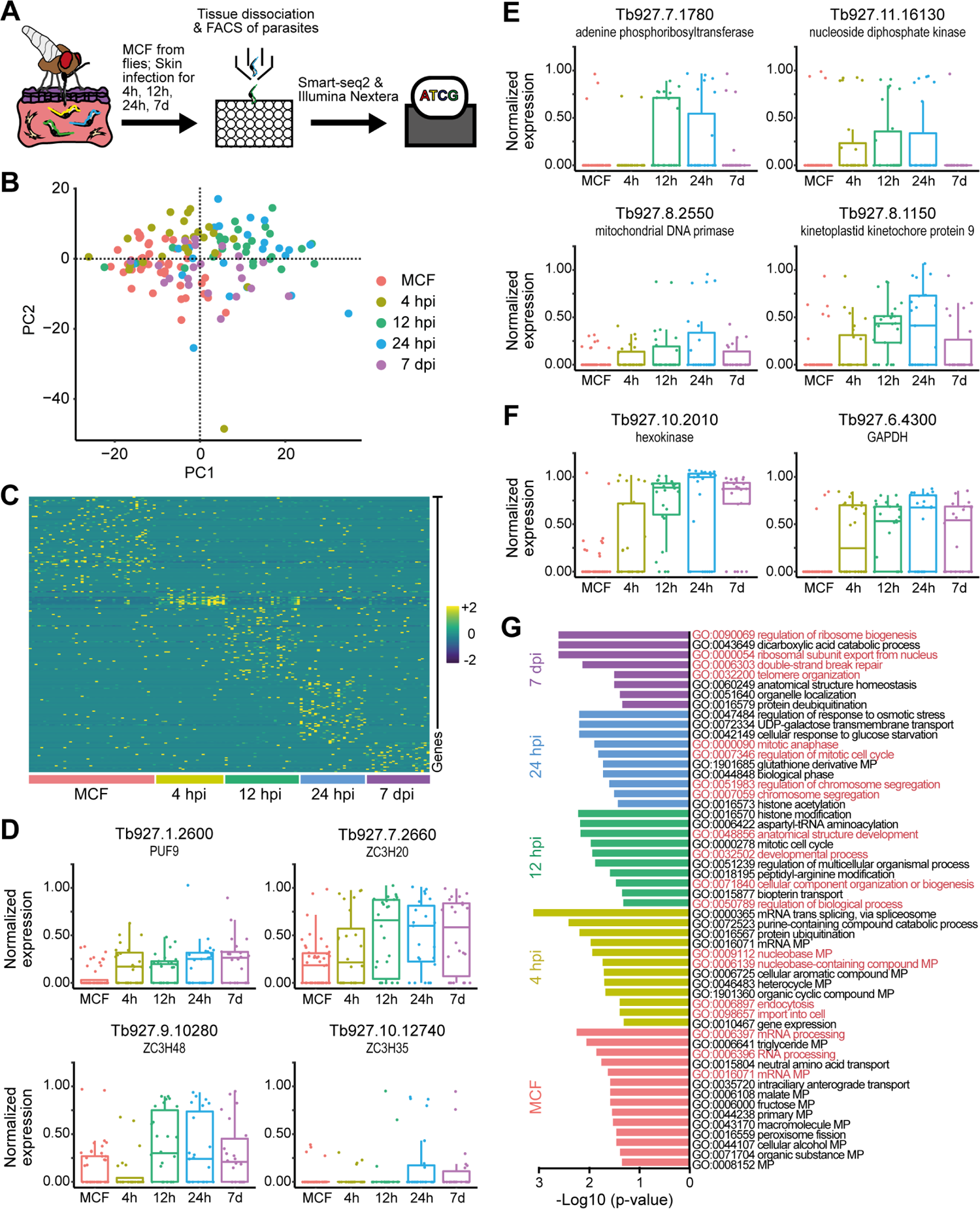
Single-parasite RNAseq reveals rapid activation of injected metacyclic trypanosomes and early events in skin infection. **(A)** Schematic representation of the experimental pipeline of single-parasite RNAseq. Tsetse-transmitted trypanosomes were isolated from infected hdSEs at 4, 12, 24 hpi, and 7 dpi. Single cells of the skin-derived parasites as well as MCFs obtained from flies were sorted into individual microwells using FACS, followed by processing with the Smart-seq2 and Nextera protocols and subsequent sequencing. **(B)** Unbiased projection of the first two principal components (PC) of the 139 trypanosome transcriptomes that passed quality control (MCF, n = 44; 4 h, n = 24; 12 h, n = 26; 24 h, n = 23; 7 d, n = 22). Each dot represents a single parasite. Colors indicate the time when the parasites were isolated from hdSEs after infection. **(C)** Heatmap showing the scaled, time-resolved expression levels of significantly upregulated genes as identified by DESeq2 (log2 fold change > 2, adjusted P value < 0.01). Each column represents a single parasite, and each row represents an individual gene. The color key from purple to cyan indicates low to high gene expression levels. **(D, E, F)** Boxplots showing the normalized expression of selected RNA-binding proteins (D), replication-associated genes (E), and glycolysis-associated genes (F) in MCFs and parasites isolated from skin at 4, 12, 24 hpi, and 7 dpi. Results are shown as median ± IQR. **(G)** Biological process-associated gene ontology (GO) terms significantly enriched in MCFs and parasites isolated from skin at 4, 12, 24 hpi, and 7 dpi. GO terms were filtered with the Revigo webtool to avoid redundancies. Bars represent the level of significance of the term enrichment.

Since different trypanosome developmental stages can reside within infected tsetse salivary glands, all transcriptomes were screened for the presence of BARP-positive parasites and three such parasites were excluded from the MCF transcriptomes (Figure S5 C). BARP-genes are specifically expressed in epimastigote forms (Urwyler et al., 2007). These parasites give rise to MCFs, but are themselves not infective. All transcriptomes were examined using an unbiased principal component analysis (PCA; Figure 5 B). Although the transcriptomes collected at different timepoints clustered closely together, the first two principal components could delineate the different timepoints. MCFs are primed for mammalian invasion and in many aspects already resemble BSFs (Christiano et al., 2017). However, based on the first principal component, MCFs (red circles) could be distinguished from parasites isolated at 12 or 24 hpi (green, blue circles, Figure 5 B).

Differential gene expression analysis revealed a number of genes specific to each timepoint - MCF: 73 genes; 4 h: 16 genes; 12 h: 61 genes; 24 h: 54 genes, and 7 d: 26 genes (Figure 5 C and Table S2). Since parasites isolated at 4 hpi had not yet started cell division (Figure 4 A) and clustered closer to MCFs (Figure 5 B), differential gene expression analysis was additionally conducted between early (MCF + 4 hpi) and late (12 hpi + 24 hpi) timepoints (Figure S5 D). Here, 24 genes were significantly upregulated at the early timepoints and 92 genes at late times (Table S2). Among the differently-expressed genes, many RNA-binding proteins (RBPs) were up or downregulated at the transcript level for all conditions. In trypanosomes, RBPs play a crucial role in the post-transcriptional regulation of gene expression and adaptation to different environments, simply because transcriptional control of Pol II-transcribed genes does not occur (Clayton, 2013, Kolev et al., 2014).

The RBPs PUF9 (Tb927.1.2600), ZC3H20 (Tb927.7.2660), ZC3H35 (Tb927.10.12740), and ZC3H48 (Tb927.9.10280) were all upregulated during skin infection (Figure 5 D). PUF9 had been previously identified as important for the progression through the cell cycle (Archer et al., 2009). In a number of publications, ZC3H20 was shown to oscillate throughout the life cycle of *T. brucei* (Christiano et al., 2017, Vigneron et al., 2020, Cayla et al., 2020, Liu et al., 2020) and to be implicated in the increase of protein translation or in mRNA stability (Erben et al., 2014, Singh et al., 2014). Thus, upregulation of PUF9 and ZC3H20 is consistent with the above results on cell cycle re-entry and protein synthesis activation. Furthermore, a rapid upregulation of replication-associated transcripts (Figure 5 E) and genes associated with glycolysis (Figure 5 F) was observed. In contrast to the fly-stages, the mammalian stages of *T. brucei* almost exclusively rely on glycolysis, as glucose is abundant in the mammalian host (Smith et al., 2017). The functions of the two RBPs ZC3H35 and ZC3H48 are not known yet, but both were described to suppress mRNAs containing *boxB* RNA hairpin-motifs in BSFs (Erben et al., 2014, Lueong et al., 2016).

More globally, Gene Ontology (GO) analysis of differently-expressed genes revealed typical signatures associated with each timepoint (Figure 5 G). While MCFs were preferentially enriched in functions associated with “mRNA processing”, parasites isolated at 4 hpi were additionally associated with GO terms such as “endocytosis”, “import into cell”, and “nucleobase metabolic processes”. Endocytosis is essential to this parasite, not only for nutrient uptake but also for survival in a hostile environment. In fact, BSFs have one of the highest rates of endocytosis of any known eukaryote (Engstler et al., 2004) and activation of endocytosis is portrayed as an adaptation to the mammalian host (Natesan et al., 2007). Parasites isolated from skin at 12 hpi were associated with GO terms reflecting reorganization of cellular structures, e.g. “anatomical structure development”, “developmental process”, or “cellular component organization or biogenesis”. Skin-derived parasites at 24 hpi were enriched in functions associated with proliferation, while parasites residing in the skin for 7 days were preferentially associated with “telomere organization”, “double-strand break repair”, and “regulation of ribosomes”. Telomeres and DNA double-strand breaks play important roles in antigenic variation and may be associated with the exchange of the metacyclic VSG coat with VSG isoforms typical of the BSF (Li, 2015). GO analysis of early (MCF + 4 hpi) and late (12 hpi + 24 hpi) timepoints resulted in a similar picture (Figure S5 E), however, with some additional early terms such as “regulation of cellular response to heat” and late terms such as “gluconeogenesis”.

Overall, the scRNAseq results confirmed that MCFs are rapidly activated in the hdSEs to give rise to a replicating parasite population. Furthermore, a variety of genes potentially involved in cell cycle activation and parasite development were identified, as well as RBPs that might control the differentiation processes.

### Full developmental competence of skin-residing parasites

It has been shown that extravascular parasites residing in the skin of infected mice can develop into the second mammalian life cycle stage, known as the stumpy form, and that skin-derived parasites can infect tsetse flies (Capewell et al., 2016). To test if the trypanosomes in the hdSE are competent to differentiate into stumpy forms, infected hdSEs were screened for the presence of parasites expressing the “Protein Associated with Differentiation 1” (PAD1), which is a stumpy-specific cell marker (Dean et al., 2009). Therefore, a trypanosome line was used harboring an NLS-GFP reporter fused to the 3’ UTR of the PAD1 gene (GFP:PAD1_UTR_). When the PAD1 gene is active, the parasites will have a GFP-positive nucleus, which is a direct indication of stumpy development (Batram et al., 2014, Zimmermann et al., 2017, Schuster et al., 2020).

On days 4 and 7 post-infection, PAD1-positive parasites were detected in the hdSEs, albeit with very low abundance (< 0.1 %; Figure 6 A and Movie 3). A screening of tsetse flies infected with skin-derived parasites at 1, 4, and 7 dpi revealed that the parasites could infect tsetse flies from 4 dpi (Figure 6 B). Detection of trypanosomes in the midgut (MG), proventriculus (PV), and salivary glands (SG) of the flies indicated that the skin-residing parasites possess, in principle, full developmental competence and can complete their entire life cycle.

**Figure 6.**
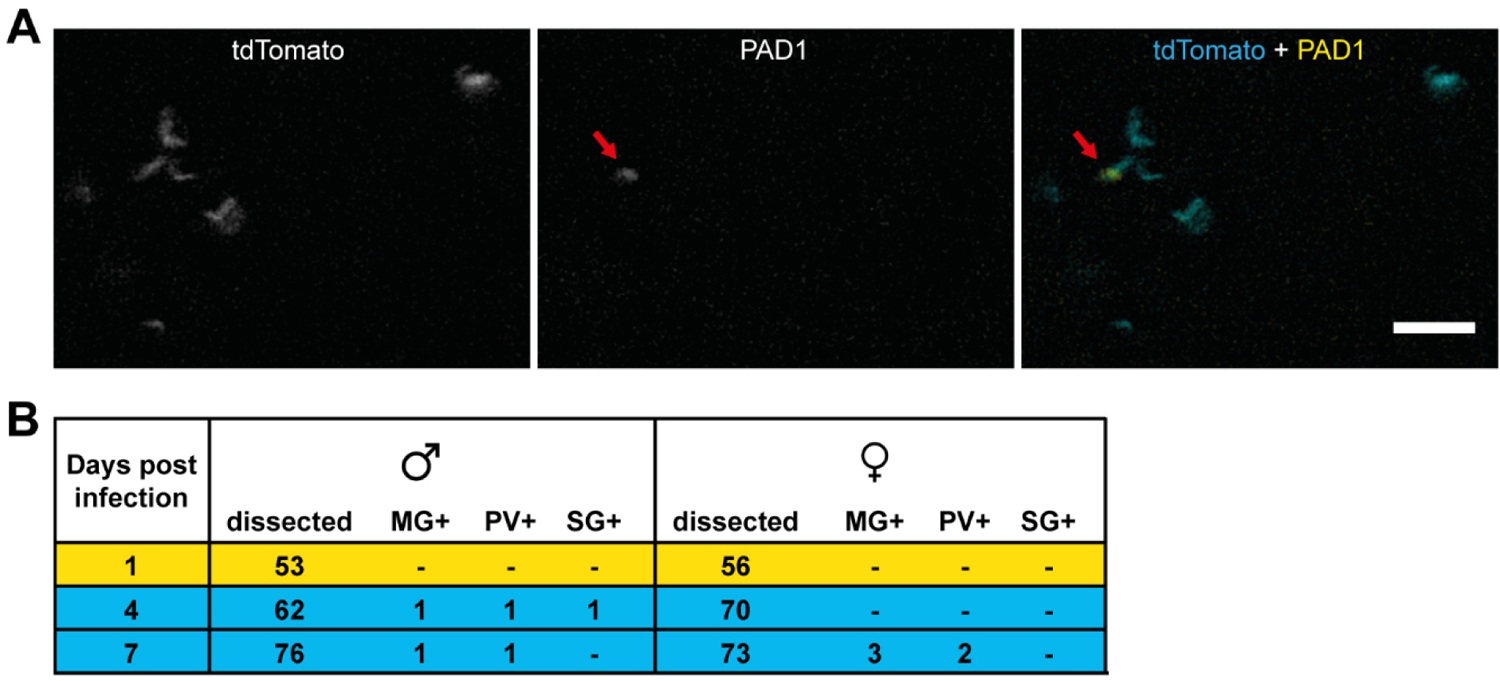
Skin-residing parasites can develop into stumpy forms and are infectious to tsetse flies. **(A)** Stereo-fluorescence microscopy of tdTomato-expressing trypanosomes in an hdSE infected for 7 days. The parasites also harbored a green fluorescent PAD1-marker fused to a nuclear localization signal as proxy for development to the stumpy stage. PAD1-positive parasites (GFP-positive nucleus, red arrow) were found in the hdSE at low incidence. Scale bar, 20 µm. **(B)** Infection of tsetse flies from hdSE-derived parasites. Trypanosomes were isolated from hdSEs at 1, 4, and 7 dpi, mixed with sheep blood, and fed to teneral tsetse flies. After 35 days, the flies were dissected and relevant organs were analyzed under a microscope and scored for the presence of trypanosomes. MG+, midgut positive, PV+, proventriculus positive, SG+, salivary glands positive.

### *T. brucei* enters a reversible quiescence program in the skin equivalents

In the second phase of skin infection (4 and 7 dpi), the trypanosomes reduced their protein synthesis rate to a basal level, which was comparable to that of cells isolated at 1 hpi, i.e. very soon after tsetse transmission (Figure 4 F and 4 G). These parasites continued to proliferate, however, albeit slowly: the population doubling time was measured at 19 h within the first 24 hpi, then slowed to 74 h within the following 3 days (day 1 to day 4) and was at 62 h in the last 3 days (day 4 to day 7), respectively (Figure 3 D, 4 A, and S4 C). In addition, replication-associated genes were downregulated at 7 dpi (Figure 5 E) and no significant increase in cell death was observed (Figure 3 E). All these findings are characteristic of quiescent cells (Rittershaus et al., 2013, Barrett et al., 2019). Thus, the parasites appeared to have launched a quiescence program. This possibility was further tested by a second independent measure of metabolic activity. The fluorescent reporter tdTomato was integrated within the rDNA locus. Expression of the fluorescent tdTomato protein therefore reported on the activity of the rDNA locus as a proxy for metabolic activity. This approach has previously been used to track quiescence in *Leishmania* (Jara et al., 2019).

In agreement with the protein synthesis measurements (Figure 4 F and 4 G), MCFs showed a significantly lower tdTomato mean fluorescence intensity (MFI) compared to cultured BSFs (Figure 7 A and S6 A). After injection, the tdTomato MFI of the skin-residing parasites initially increased within the first 24 h, but then decreased by 2.6- and 4-fold on days 4 and 7 post-infection, respectively. To test whether this phenotype could be reversed, the migration from the skin to the bloodstream was mimicked by extracting skin-residing parasites on day 7 and transferring them into culture medium supplemented with methylcellulose to mimic blood viscosity. A significantly increased protein synthesis rate (Figure 4 F and 4 G) and tdTomato MFI (Figure 7 A and S6 A) was detected within 24 h after skin-to-medium transfer. Moreover, the tdTomato MFI was stable for at least 3 days after transfer (Figure 7 A). Upon transfer to culture medium, the population doubling time dropped to 7.2 h (Figure 7 B), a value comparable to the 6.2 h of the BSF wildtype strain (Figure S2 A).

**Figure 7.**
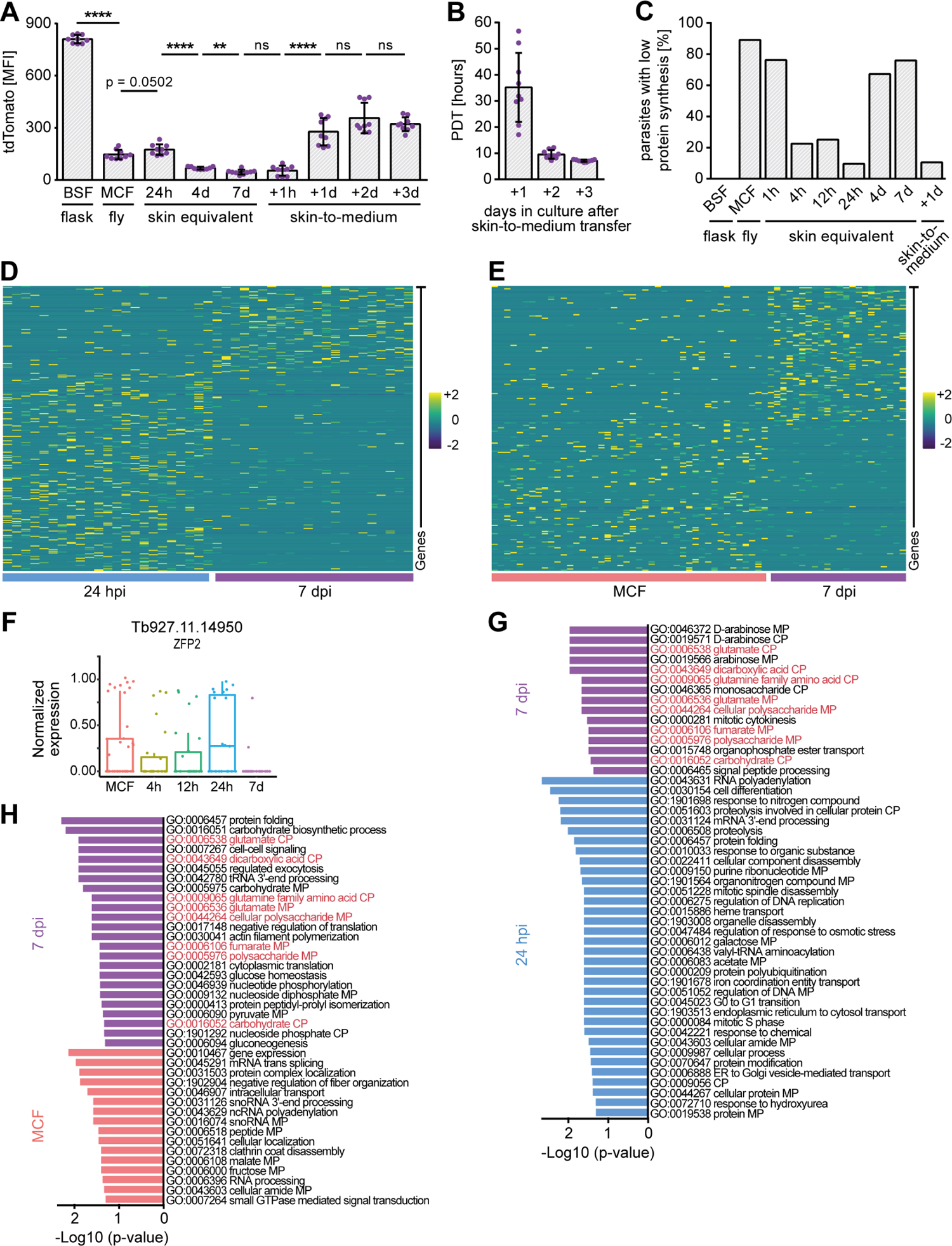
Trypanosomes reversibly enter a quiescent state in the skin that is characterized by a unique gene expression profile. **(A)** Flow cytometry quantification of the tdTomato MFI of skin-resident trypanosomes compared to BSFs, MCFs, and skin-parasites at 7 dpi, which were transferred to culture medium for up to 3 days (skin-to-medium). The tdTomato reporter is located in the rDNA locus, and thus a proxy for the metabolic state of the parasite. Graphs represent mean ± SD (n = 9). Unpaired t-test, **** p < 0.0001, ** p < 0.01, ns, not significant. **(B)** Parasites were isolated from hdSEs at 7 dpi and transferred to HMI9 culture medium supplemented with methylcellulose to mimic blood viscosity. The PDTs were recorded for up to 3 days. Data represent mean ± SD (n = 9). **(C)** Quantification of cells classified as “parasites with low protein synthesis”, indicated by the red line in Figure 4 G, in BSFs, MCFs, skin-residing trypanosomes, and parasites at 7 dpi which were transferred to culture medium for 1 day (skin-to-medium) **(D, E)** Heatmaps showing the scaled expression levels of upregulated genes of MCFs and skin-residing parasites at 24 hpi and 7 dpi, as identified by DESeq2 (log2 fold change > 2, adjusted P value < 0.01). Each column represents a single cell, each row represents an individual gene. The color key from purple to cyan indicates low to high gene expression levels. **(F)** Boxplot showing the normalized expression of the RNA-binding protein ZFP2 in MCFs and parasites isolated from skin at 4, 12, 24 hpi, and 7 dpi. Results are shown as median ± IQR. **(G, H)** Biological process-associated GO terms significantly enriched in MCFs and parasites isolated from skin at 24 hpi and 7 dpi. GO terms were filtered with the Revigo webtool to avoid redundancies. Bars represent the level of significance of the term enrichment.

Since protein synthesis is the most energy-consuming cellular process, it is downregulated to a minimal level during quiescence (Hofmann and Hand, 1994). Thus, the percentage of skin-residing parasites with a low protein synthesis rate was quantified for each timepoint. The lowest synthesis rate measured in cultured BSFs was used as a threshold value (Figure 4 G, red line). At 4 and 7 dpi, 67 % and 76 % of skin-residing parasites revealed a strongly reduced protein synthesis rate. Transfer to culture medium reversed this phenotype and the number of parasites with low protein synthesis decreased to 10.5 % within 24 h (Figure 7 C), and the protein synthesis rate was comparable to cultured BSFs (Figure 4 G).

To test whether the induction of a quiescent state can be attributed to the skin microenvironment, metacyclic trypanosomes harvested from tsetse flies were inoculated directly into culture medium supplemented with methylcellulose (Figure S6 B). Within the observation period of 10 days, the parasites entered exponential growth and had a population doubling time of 6.8 h. This is comparable to the 7.2 h of the extracted skin-residing parasites that were transferred into culture medium (Figure 7 B). However, no reduction in parasite growth was observed over the 10-day period, which could indicate an entry into quiescence. Interestingly, this proliferative trypanosome population was 14.9 % AnTat 1.1-positive after 7 days (Figure S6 C), in strong contrast to the < 0.1 % of AnTat 1.1-positive skin-dwelling parasites observed on days 4 and 7 (Figure 4 E).

To investigate the quiescent trypanosome population in more detail, differential gene expression analysis between parasites isolated from infected skin 24 hpi and 7 dpi was performed (Figure 7 D). The comparative scRNAseq analysis of highly active (24 hpi) and inactive (7 dpi) parasites identified 226 genes that were significantly upregulated at 24 hpi and 99 genes specifically upregulated at 7 dpi (Table S3). In addition, the transcriptomes of MCFs were compared with parasites isolated at 7 dpi, as both forms possess a reduced metabolism (Figure 7 E). Here, 127 genes enriched in MCFs and 115 genes specific for 7 dpi-parasites were retrieved (Table S3). Among the differentially-expressed genes, the RNA-binding protein ZFP2 (Tb927.11.14950) was downregulated at 7 dpi (Figure 7 F) and could be one factor that regulates the development of a quiescent trypanosome form in the skin.

GO enrichment analysis revealed several metabolic and catabolic processes that were regulated differentially between parasites isolated at 24 hpi and 7 dpi, particularly with regard to arabinose (Figure 7 G). Compared to MCFs, 7 dpi-parasites were enriched in functions associated with “pyruvate metabolic process”, “glucose homeostasis”, or “protein folding” (Figure 7 H). GO terms that occurred in both analyses for 7 dpi-parasites were functions associated with amino acid (glutamate, glutamine), polysaccharide, and dicarboxylic acid metabolism (fumarate). The underlying gene of the GO terms associated with glutamate is a glutamate dehydrogenase (Tb927.9.5900), and it has been shown that glutamate dehydrogenase is induced in quiescent yeast and epithelial cells (Brauer et al., 2005, Murphy et al., 2015, Coloff et al., 2016).

In conclusion, we have developed a novel skin tissue model that resembles native human skin in key anatomical, cellular, and functional aspects and can be infected with trypanosomes through the bite of the tsetse fly. We have shown that the parasites almost immediately reactivate the cell cycle and establish a proliferative skin tissue population. In the second phase of skin residence, the trypanosome population shared common features described for quiescent cells, namely slow growth, reduced DNA synthesis, and downregulation of protein translation. In addition, their metabolism and transcriptome differed from the highly proliferative trypanosomes in the skin, and they could be reactivated and returned to an active state. The results demonstrate the existence of a quiescence program without cell cycle-arrest in trypanosomes, which may well represent an adaptation to the skin as niche for parasite persistence.

## Discussion

We have established an advanced skin model and validated it through infection with trypanosomes. By choosing the natural transmission route, the bite of an infected tsetse fly, we have been able to detail the early events of trypanosome infection with unprecedented resolution, and we clearly show that the natural host environment is required for parasite development.

Tsetse-transmitted cell cycle-arrested MCFs are rapidly activated in the artificial skin and establish a proliferative trypanosome population. This process is accompanied by (I) reactivation of protein synthesis, (II) re-entry into the cell cycle, (III) acquisition of a BSF-like morphology, and (IV) increased motility.

We found that the parasites re-entered the cell cycle already between 6 and 12 hours. Moreover, protein synthesis is reactivated very rapidly, after just 1 hour, and peaks within 1 day. At that time the rate resembles that of *in vitro* cultured BSFs. These findings are consistent with natural infections of mice showing that MCFs start multiplying in the skin within 18 hours of transmission (Caljon et al., 2016). Single-parasite RNAseq revealed an upregulation of two RNA-binding proteins (RBPs), PUF9 and ZC3H20, within 4 hours. These proteins have been shown to be involved in replicative and translational processes (Archer et al., 2009, Erben et al., 2014). In addition, we found upregulation of two previously undescribed RBPs, ZC3H35 and ZC3H48, which could well be functional during differentiation of the parasites in the skin. In contrast to the fly-stages, which use proline to feed their tricarboxylic acid (TCA) cycle, glucose is the only carbon source for ATP production described for replicative BSFs (Smith et al., 2017). MCFs pre-adapt by downregulation of mRNAs for components of the TCA cycle, but retain their protein counterparts. In addition, enzymes of the glycolytic pathway are upregulated at the protein level, but not at the transcript level (Christiano et al., 2017). Supporting this, we show that genes associated with glycolysis were rapidly upregulated within 4 hours post-infection in the skin. The morphology of MCFs is similar to that of BSFs, but their cell body and the free distal part of their flagellum are shorter (Vickerman, 1962). After 1 day, we observed that the skin-residing parasites exhibited a BSF-like morphology, accompanied by an increased swimming speed. This is consistent with observations that BSFs have a higher maximum velocity and flagellar beat frequency compared to MCFs (Bargul et al., 2016, Schuster et al., 2017, Kruger et al., 2018). An increased swimming speed could influence the spread of the parasites in the dermis and entry into the draining lymph, where parasites can be detected 1 - 2 days after infection (Barry and Emery, 1984). The exchange of the VSG coat is a known hallmark of the MCF to BSF transition (Barry et al., 1998) and the switch from expression of metacyclic to bloodstream form VSGs occurs between days 4 and 6 post-infection (Esser and Schoenbechler, 1985). The VSG repertoire of the pleomorphic strain EATRO 1125 used in this study is not known and predicting which BSF VSG isoforms are expressed is not possible. The only VSG known to be expressed in EATRO 1125 is the VSG AnTat 1.1. Thus, the stochastic reactivation of this one specific BSF expression site should result in a subpopulation of 5 - 7 % of trypanosomes displaying VSG AnTat 1.1 (provided that the strain harbors 20 or 15 expression sites, respectively). However, positive immunostaining for this BSF VSG isoform was only detected in < 0.1 % of the skin-residing parasites 4 and 7 days post-infection and contrasts with the observation that MCFs differentiated in suspension culture had a proportion of 14.9 % of AnTat 1.1-positive parasites. A similar observation has been made with parasites of the same strain residing in adipose tissue. Here, the active VSG AnTat 1.1 was 3-fold downregulated compared to parasites in the bloodstream (Trindade et al., 2016). These observations suggest that trypanosome populations residing in different niches of the same host (e.g., skin, adipose tissue, blood) express different VSGs. Furthermore, it strengthens our finding that the parasites in the skin at day 7 are not *bona fide* BSFs.

Although the reason(s) why trypanosomes accumulate and persist in mammalian skin remain unknown (Caljon et al., 2016, Capewell et al., 2016), we clearly show that the skin environment strongly influences the parasites. After re-entry into the cell cycle and an initial phase of proliferation, the skin-residing trypanosomes slow down growth, reduce DNA synthesis, and downregulate protein translation and metabolism. Concomitantly, they have very low rates of parasite death. All these characteristics are hallmarks of quiescent cells (Coller, 2011, Daignan-Fornier and Sagot, 2011, Laporte et al., 2011, Rittershaus et al., 2013, Barrett et al., 2019). We postulate that the initial phase of trypanosome proliferation functions in establishing a robust population in the host skin. Once this is achieved a second adaptation step yields a slow growing, skin-residing trypanosome population, which contributes to parasite persistence. Interestingly, the trypanosomes do not actively leave the skin model, as very few parasites were found in the surrounding culture medium. Furthermore, when the parasites were extracted from the skin and transferred to culture medium, they resumed rapid growth. Differential gene expression analysis on the single-cell level revealed that the quiescent parasites clearly differ from the early proliferative trypanosomes after infection. All the above prompts us to propose that trypanosomes transmitted by the tsetse fly into skin enter a persister-like state, the quiescent skin tissue form (STF).

The quiescent STFs are distinct from the two other known quiescent life cycle stages of *T. brucei*, the metacyclic and the stumpy stage, and may represent an adaptive response to the skin microenvironment. First, MCFs and stumpy forms are cell cycle-arrested and require a host change for their reactivation (Szoor et al., 2020). In contrast, the STFs described here replicated very slowly and can be reactivated within the same host, namely when they transit from skin via lymphatic vessels to the bloodstream. Second, the STFs did not, or only very rarely, differentiate to the cell cycle-arrested stumpy stage. The quiescent parasite population in the skin did not express the stumpy-specific marker protein PAD1. Third, the transcriptome of STFs clearly differed from MCFs. Fourth, the morphology of STFs resembled that of BSFs, and fifth, STFs swam faster in the skin than MCFs. Theoretically, induction of quiescence by macro- or micro-environmental conditions can be distinguished from quiescence coupled to differentiation. Moreover, quiescence can also be induced when conditions are suitable to sustain proliferation, and entry and exit into quiescence is independent of the G1 phase of the cell cycle (Laporte et al., 2011, Daignan-Fornier and Sagot, 2011, Fiore et al., 2018, Barrett et al., 2019, Sagot and Laporte, 2019).

We do not consider the STFs as an as yet undiscovered life cycle stage of *T. brucei*, as these forms can be bypassed by direct inoculation of metacyclic parasites into the host’s body cavity or blood. We rather suggest that the formation of STFs (and probably also the adipose tissue forms (ATFs) residing in fat (Trindade et al., 2016)) are evolutionary examples for the amazing flexibility in parasite life cycles. Trypanosomes are prime examples for this, because they thrive in various host animals and in body fluids with very different physicochemical properties. In adipose tissue, trypanosomes have been shown to metabolically adapt as ATFs (Trindade et al., 2016). ATFs have a BSF-like morphology but, as a functional adaptation to adipose tissue, can utilize fatty acids as a carbon source, unlike BSFs. However, it remains unclear whether ATFs similarly exhibit reduced metabolism and slow growth. The ability to flexibly and specifically respond to environmental cues is an as yet underestimated hallmark of parasitism.

We found that the STFs replicate 10-times slower and have a protein synthesis rate 3.5-times lower than cultured BSFs. Similar observations were made with *Mycobacterium tuberculosis*, which has a 5-fold higher doubling time during persistence (Munoz-Elias et al., 2005, Gill et al., 2009), and quiescence in *Saccharomyces cerevisiae* is accompanied by a 20-fold decrease in protein synthesis (Fuge et al., 1994). Slow growing persister cells have been described for a few other protist parasites, such as *Leishmania* or *Trypanosoma cruzi*, but not for African trypanosomes (Barrett et al., 2019). In the related protist, *Leishmania mexicana*, quiescent amastigotes in murine lesions show very slow growth with a doubling time of 12 days and low rates of protein turnover (Kloehn et al., 2015), and slow-growing intracellular amastigotes of *Leishmania major* with a doubling time of 60 hours have also been reported (Mandell and Beverley, 2017). In *Trypanosoma cruzi*, chronic infections are associated with a reduced replication rate of the parasite in the colon (Ward et al., 2020).

Skin-residing trypanosomes in a mouse model can develop into stumpy forms and their proportion has been determined to be 20 % on day 11 post-infection (Capewell et al., 2016). This is actually rather late. Although we found stumpy forms in our skin model, their proportion was very low with < 0.1 % of all parasites on days 4 and 7. On one hand, this shows that the trypanosomes in the skin model are, in principle, capable of stumpy formation at these early timepoints. On the other hand, the STFs should be resistant to stumpy induction, as the postulated paracrine induction of stumpy formation generated by trypanosome excreted oligopeptidases (Rojas et al., 2019) would have yielded a much larger proportion of stumpy parasites in the confined skin environment. Similarly, it has been shown that quiescent myoblasts and fibroblasts are resistant to terminal differentiation (Coller et al., 2006). Alternatively, STFs do not secrete oligopeptidases. The hypothesis of specific skin tissue forms is further supported by the very low infection rates of tsetse flies that were fed with skin-derived trypanosomes. We have recently shown that a single bloodstream stage trypanosome suffices to successfully infect a tsetse fly, and that it does not matter if this is a proliferative BSF or stumpy stage parasite (Schuster et al., 2020). Thus, if the STFs would be BSFs, we would expect much higher fly infection rates. It is important to note that the same pleomorphic trypanosome serodeme AnTat 1.1 was also used in the tsetse infection study. This suggests that STFs are not tsetse-infective.

In conclusion, our experiments are compatible with the following scenario. The tsetse fly deposits cell cycle-arrested MCFs into the host skin. The feeding insect ruptures blood vessels. MCF parasites that directly enter the circulation through the pooled blood, develop to the proliferating BSF stage. Parasites that do not enter the bloodstream but stay in the skin start proliferating in tissue spaces, and in this way establishing a viable population of proliferative trypanosomes within 1 day. At this timepoint they adapt to the skin environment as quiescent STFs, which can persist in the host’s skin for extended periods. As the skin interstitial fluid is continuous with the lymphatic system, STF parasites will eventually be drained by suction forces into the lymph, where they start proliferating as BSFs. The skin acts as a reservoir for African trypanosomes (Capewell et al., 2016, Camara et al., 2020) and the hidden, quiescent, but motile STFs could act as a source of parasites that can continuously repopulate the blood.

Interestingly, after the initial immune response of the host against the injected metacyclic trypanosomes in the skin, the skin-dwelling parasites appear not to trigger any further major inflammatory cell infiltration (Capewell et al., 2016, Mabille and Caljon, 2020). The underlying host and parasite factors are unknown. Our discovery of STFs in a close to nature, tractable tissue model might provide an experimental system for tackling the important question as to how the parasites might be slipping under the radar of the inflammation response. In recent years, there has been increasing evidence that trypanosomes persist in the skin of aparasitemic and asymptomatic individuals (Capewell et al., 2016, Camara et al., 2020). It has also been suggested that these individuals contribute to the transmission of the disease (Koffi et al., 2006, Kabore et al., 2011, Capewell et al., 2019). In addition, case reports of longstanding infections with little to no symptoms (Ilboudo et al., 2011, Jamonneau et al., 2012, Sudarshi et al., 2014, Berthier et al., 2016) suggest that at least some individuals may develop latent trypanosome infections, in which quiescent, slowly replicating trypanosomes residing in the skin may play an important role. Furthermore, the skin quiescence might explain some of the treatment failures in humans (Richardson et al., 2016). Quiescent cells are usually less susceptible to chemotherapeutic drugs (Witkowski et al., 2010).

Lastly, we would like to emphasize that our work introduces the host-parasite interface as a useful tool for the qualification of human tissue models. The advanced skin tissue model developed in the course of this study is permissive for dual scRNAseq studies of skin cells and parasites during early stages of a natural, vector-borne infection. The in-depth analyses described here revealed the existence of skin tissue forms of *T. brucei* and, at the same time, proved that our skin model delivers all cues required for induction of parasite development. In contrast to laboratory animals, tissue models allow the use of primary human cells and they can be iteratively advanced. The introduction of new cell types, such as macrophages will enable infection studies with other important parasites, for example *Leishmania*.

## Author Contributions

Conceptualization, C.R., A.-E.S., F.G.-B., and M.E.; Methodology, C.R., F.I., L.H., P.F. and T.F.; Formal Analysis, C.R. and E.V.; Investigation, C.R., F.I., L.H. and P.F.; Writing - Original Draft, C.R. and M.E.; Writing - Review & Editing, all authors; Supervision, H.W., A.-E.S., F.G.-B., and M.E.; Funding Acquisition, M.E.

## Acknowledgments

The authors thank Panagiota Arampatzi, Nina DiFabion, Tobias Krammer, Christophe Toussaint, and Oliver Dietrich for guidance and useful discussions regarding scRNAseq; Stephan Löwe for his help in producing the fly video; Elisabeth Meyer-Natus for guidance and support with scanning electron microscopy; Brooke Morriswood for critical reading of the manuscript and providing helpful and critical feedback; Kathrin Weißenberg for logistical help in isolating trypanosomes from skin for scRNAseq; Tim Krüger for guidance on parasite tracking in skin and data analysis. C.R. is member of the Graduate School of Life Sciences in Würzburg and is funded by the DFG GRK 2157 (3D Tissue Models to Study Microbial Infections by Obligate Human Pathogens). M.E. is further supported by DFG grants EN305, SPP1726 (Microswimmers - From Single Particle Motion to Collective Behaviour), GIF grant I-473-416.13/2018 (Effect of extracellular *Trypanosoma brucei* vesicles on collective and social parasite motility and development in the tsetse fly), the EU ITN Physics of Motility, and the BMBF NUM Organo-Strat. M.E. is a member of the Wilhelm Conrad Roentgen Center for Complex Material Systems (RCCM).

## Materials and Methods

### Contact for Reagent and Resource Sharing

Further information and requests for resources and reagents should be directed to and will be fulfilled by the Lead Contact, Markus Engstler

### Experimental Model and Subject Details

#### Ethical clearance statement

Normal primary human epidermal keratinocytes (NHEK) and dermal fibroblasts (NHDF) were isolated from biopsies of preputial skin from juvenile donors aged between 2 and 5 years. All donors’ legal representative(s) provided full informed consent in writing. The study was approved by the local ethical board of the University of Würzburg (vote 182/10 and 280/18-SC).

#### Isolation of primary human cells and culture

Biopsies of preputial skin were washed with PBS containing Mg^2+^/Ca^2+^ and after removal of excess fat and connective tissue, cut into 3 mm x 10 mm sized pieces. Subsequently, pieces were incubated with 2 U/ml dispase at 4 °C overnight and the epidermis was separated from the dermis with forceps. To isolate NHEK, epidermal pieces were minced and incubated with 0.05 % trypsin/EDTA at 37 °C for 5 min, followed by thorough resuspension to release cells. A 100 µm cell strainer was used to filter cell suspension and keratinocytes were cultured in EpiLife basal medium supplemented with 1 % human keratinocyte growth supplement (HKGS) and 1 % penicillin/streptomycin. To isolate NHDF, dermal pieces were minced and incubated with 500 U/ml collagenase type XI at 37 °C for 45 min. Dermal pieces were washed with DMEM supplemented with 10 % fetal calf serum (FCS), 1 % non-essential amino acids (NEAA), and 1 % penicillin/streptomycin and transferred into cell culture flasks. Dermal pieces were removed after 5 days after fibroblasts had grown out.

For both cell types, medium was changed every 2 - 3 days and cells were passaged when reaching 70 - 90 % confluency by Accutase-(NHEK) or Trypsin-treatment (NHDF). To generate dermal equivalents or high-density skin equivalents cells from passage 3 to 5 were used.

#### High-density primary human skin equivalent

The first step in the generation of high-density primary human skin equivalents (hdSEs) involves the production of the high-density dermal equivalent (hdDE). The compression reactor was assembled and the membrane of 12 Snapwell™ inserts was perforated with 42 microneedles to improve permeability before the inserts were placed in the designated slots in the reactor. Next, 800 µl of reconstituted collagen solution (c_collagen_ = 6.7 mg/ml) containing 45,200 NHDF was filled in each of the 12 compression chambers. Briefly, 8 ml of cold 10 mg/ml collagen type I from rat tail dissolved in 0.1 % acetic acid was taken up in a 20 ml syringe without air bubbles using a sterile 13G cannula. Subsequently, 6.78 x 10^5^ NHDF were resuspended in 4 ml of cold neutralization solution (2x DMEM, 3 % FCS, 3 % 3M HEPES, 1 % chondroitin sulfate, pH = 8.5) and the suspension was taken up in a 20 ml syringe using a sterile 18G cannula. Next, both syringes were connected to a three-way valve and mixed by alternately pressing the two syringes. After 8 - 10 mixing steps, the empty syringe was disconnected and replaced by a Safeflow® needlefree injection port. A 10 ml Combitip® connected to an Eppendorf Multipette M4 was inserted into the injection port and by pressing the syringe the neutralized NHDF-containing collagen solution was transferred into the Combitip®. 800 µl were dispensed air bubble-free in each compression chamber and to ensure complete gelation of collagen, the reactor was incubated 30 min under a sterile workbench and an additional 30 min in a cell culture incubator at 37 °C and 5 % CO_2_.

The lid of the reactor was attached to the linear motor and the solidified collagen gels were compressed by a factor of 7 by moving the lid downwards at constant 2 x 10^-6^ m/s until the reactor was completely closed. The reactor was opened and compressed air was introduced via the adapter into the lid and the compression punches to release the compressed dermal equivalents without damage. Each hdDE had a calculated collagen concentration of 46.9 mg/ml and was characterized by a diameter of 12 mm and a height of 1 mm, resulting in a volume of 113 µl. hdDEs were cultured with DMEM supplemented with 10 % FCS, 1 % NEAA, and 1 % penicillin/streptomycin.

On day 3, 250 µl of EpiLife basal medium supplemented with 1 % HKGS, 1 % penicillin/streptomycin, and 1.44 mM CaCl_2_ (= E2 medium) containing 5 x 10^5^ NHEK were added to the apical side of each hdDE to generate the epidermis of the hdSE. The next day, the E2 medium was aspirated and air-liquid interface (ALI) culture was initiated with E2 medium supplemented with 10 ng/ml human keratinocyte growth factor (hKGF) and 252 µM L-Ascorbic acid 2-phosphate (= E3 medium). ALI culture was continued until day 23 at 37 °C and 5 % CO_2_ with medium change every 2 - 3 days.

#### Trypanosome strain and culture

The pleomorphic *Trypanosoma brucei* EATRO 1125 AnTat 1.1 strain harboring the two reporters *GFP::PAD1_UTR_* and *tdTomato* was used for all experiments. The construct *GFP::PAD1_UTR_* is a reporter for the fly-transmissible stumpy form, in which the green fluorescent protein GFP is coupled to the 3’UTR of the stumpy marker PAD1 (Dean et al., 2009). The GFP protein is targeted into the nucleus via a nuclear localization signal (NLS). The fluorescent protein tdTomato is located in the ribosomal DNA locus (Xong et al., 1998) and expressed in all lifecycle stages of the parasite. Bloodstream forms were cultured in suspension below 5 x 10^5^ cells/ml in viscous HMI9 medium supplemented with 10 % FCS and 1.1 % methylcellulose (Vassella et al., 2001) at 37 °C and 5 % CO_2_.

#### Tsetse fly colony

Tsetse flies of the subspecies *Glossina morsitans morsitans* were kept in an insectary at 27 °C and a relative humidity of 70 % and fed three times a week through a silicon membrane with preheated defibrinated sterile sheep blood.

### Method Details

#### Compression reactor design and fabrication

The computer-aided design (CAD) of the compression reactor was conducted with SolidWorks® and the reactor was manufactured from polyether ether ketone (PEEK). The reactor allowed simultaneous compression of 12 collagen gels in Snapwell™ inserts. A perforated plate beneath the insert holder prevented the insert membranes from rupturing during compression and a liquid collecting tray collected the displaced water from the gels. The compression chamber increased the compression volume per insert up to 2.8 ml, and all parts could be firmly connected to the compression chamber by tightening the swing screws located on both sides of the liquid collecting tray.

The 12 cylindrical compression punches of the lid were designed to fit exactly into the 12 openings of the compression chamber with a minimum of friction. Due to their defined length the distance between the insert membranes and the lower edge of the compression punches was always exactly 1 mm after complete compression. An adapter for the attachment to the linear motor with a compressed air connection could be mounted centrally on the lid via a screw thread. Via the connection compressed air with a pressure of 4 - 6 bar could be introduced and was distributed through the sealed cavity in the lid into the 0.5 mm diameter channels, which pass centrally through each of the 12 compression punches. The compressed air was needed to release the compressed collagen gels in the Snapwell™ inserts from the compression punches without damage.

Compression was performed with a linear motor from the company NTI AG LinMot & MagSpring (Switzerland). The linear motor was attached to a stainless steel motor mount via a motor flange and connected to a computer via a servo drive. The motor was controlled using the LinMot®-Talk software.

#### Skin dissociation and quantification of cells

To isolate NHDF and trypanosomes from skin equivalents, the epidermis was first removed from the dermis with forceps. The dermis was cut into small pieces with a scalpel and incubated in 1 ml of trypanosome dilution buffer (TDB; 5 mM KCl, 80 mM NaCl, 1 mM MgSO_4_, 20 mM Na_2_HPO_4_, 2 mM NaH_2_PO_4_, 20 mM glucose, pH = 7.6) containing 250 µg Liberase TL in a water bath at 37 °C for 45 min. Cells were collected by centrifugation at 2000 g for 2:30 min, resuspended in TDB, and counted.

#### Flow cytometry

Skin equivalents were dissociated and the cells were resuspended in 500 µl TDB, then filtered through a 35 µm cell strainer. 20 min and 5 min before analysis, 2 µM Calcein AM Violet and 2 µM NuclearGreen were added, respectively. Samples were then processed on a FACSAria III cytometer (BD Biosciences). Trypanosomes were analyzed by gating first on the tdTomato signal to exclude NHDF and extracellular matrix proteins from analysis. Next, viable parasites were selected by setting a gate on the Calcein AM signal and finally, cell cycle distribution was determined by NuclearGreen fluorescence. Cell size of parasites was determined by using the width signal of the forward scatter (FSC-W). Analysis was performed using CellQuest Pro™ software (BD Biosciences).

#### Infection of tsetse flies

Tsetse flies were routinely infected with stumpy stage trypanosomes within 72 hours post-eclosion. BSFs were grown to a density of 5 x 10^5^ cells/ml and cultured for another 48 hours without dilution to induce stumpy formation by density. Stumpy forms were harvested by diluting cultures 1:4 with TDB and subsequent filtration to remove methylcellulose, followed by centrifugation at 1400 g for 15 min at 37 °C. Parasites were resuspended in sheep blood at a concentration of 4 - 8 x 10^6^ parasites/ml, supplemented with 60 mM N-acetyl-D-glucosamine and 12.5 mM glutathione, and fed to flies. Positive flies were selected by a salivation test (mature salivary gland infection) after 5 weeks.

To infect flies with skin-derived trypanosomes, infected skin equivalents were dissociated and viable parasites were sorted based on their tdTomato and Calcein AM signal with a FACSAria III cytometer (BD Biosciences) to exclude NHDF and extracellular matrix proteins. Parasites were resuspended in sheep blood at 5 x 10^4^ cells/ml and fed to teneral tsetse flies with their first blood meal. Flies were dissected after 5 weeks and screened for trypanosomes present in the midgut, proventriculus, and salivary glands.

#### Natural infection of skin and culture

On day 15 of skin equivalent culture, the E3 medium was exchanged with infection medium (INF; 1:1 mixture of E3 and HMI9 medium supplemented with 1 % Anti/Anti, 1 % penicillin/streptomycin, and 440 µM CaCl_2_). Skin equivalents aged between 16 and 23 days, depending on further experiments and duration of culture, were removed from the Snapwell™ inserts. Three skin equivalents were stacked on top of each other on an aseptic microscope slide on a heating plate set to 37 °C and 6 tsetse flies with a mature salivary gland infection (SG+) were allowed to bite into the stack. In order to have sufficient quantities and equal numbers of parasites for downstream analysis per skin equivalent, the positions were changed and another round of 6 SG+ flies were allowed to bite into the stack. This was repeated 3 times, so that each of the three skin equivalents was once on each position (bottom, middle, top). Stacking was necessary because the proboscis of the tsetse fly has a length of 2 mm, whereas the hdSEs have a standard height of 1 mm. Without stacking, the flies simply bit through the hdSEs and deposited most of the parasites underneath. Infected skin equivalents were cultured on sterile Whatmann® paper at the air-liquid interface with 7 ml of INF medium at 37 °C and 5 % CO_2_. Medium was changed every 2 - 3 days.

#### Assessment of injection depth

Cell-free high-density dermal equivalents were used since no further culture was carried out. Three equivalents were stacked on top of each other and 18 SG+ tsetse flies were allowed to bite into the stack without rotation. Immediately after infection, dermal equivalents were dissociated and trypanosomes were counted by flow cytometry based on their tdTomato signal. Since each high-density dermal equivalent had a standardized height of 1 mm, a depth of up to 3 mm could be simulated with a resolution of 1 mm.

#### Assessment of protein synthesis rate

Skin equivalents were dissociated and trypanosomes were resuspended in 500 µl of pre-warmed methionine-free RPMI containing 10 % FCS, 1 % Anti/Anti, 1 % penicillin/streptomycin, and either 50 µM L-Azidohomoalanine (AHA) or 50 µM L-methionine as negative control. As a further control, BSFs from cell culture flasks were included in each experiment. All samples were incubated in a cell culture incubator for 1 hour at 37 °C. Subsequently, cells were centrifuged at 2000 g for 2:30 min, resuspended in 500 µl TDB supplemented with 2 µM Calcein AM Violet and filtered through a 35 µm cell strainer.

Viable parasites were sorted based on their tdTomato and Calcein AM signal with a FACSAria III cytometer (BD Biosciences) to exclude NHDF and extracellular matrix proteins. 1 - 5 x 10^4^ parasites were sorted in 24-well plates containing coverslips pre-coated with poly-L-lysine (Ø 15 mm) and filled with 500 µl 3 % PFA (125 µl TDB + 375 µl 4 % PFA). 24-well plates were centrifuged at 1500 g for 5 min to attach parasites to the coverslips and washed once with 500 µl PBS containing 3 % BSA (PBS-B). Trypanosomes were permeabilized with 500 µl PBS-B containing 0.5 % Triton X-100 for 20 min and washed twice with 500 µl PBS-B. 300 µl of Click-iT reaction cocktail (prepared according to the manufacturers’ instructions) were added to each cover slip and incubated for 30 min at room temperature (RT) protected from light. Following a further washing step with 500 µl PBS-B, the coverslips were carefully taken out of the cavities of the 24-well plate and mounted upside down on a microscope slide with 7.5 µl Fluoromount-G with DAPI.

At least 136 trypanosomes from three individual experiments were imaged per timepoint with a DMI6000B wide-field fluorescence microscope (Leica), equipped with a 100x oil objective (NA 1.4) and a DFC365FX camera (pixel size 6.45 µm). Images were taken by focusing in the DAPI channel on the cell nucleus and kinetoplast. Subsequently, images were taken in brightfield, DAPI and AHA channels. Images were then analyzed with the ImageJ/Fiji software. To quantify the fluorescence intensity of incorporated AHA or L-methionine, a rectangular region of interest (ROI) was drawn around individual parasites. The area, mean fluorescent intensity, and integrated density of the ROI was measured in the AHA channel along with several adjacent background measurements using the built-in measurement program (Analyze > Measure). The corrected total cell fluorescence (CTCF) of individual cells incubated with either AHA or L-methionine was calculated using the following formula: CTCF = integrated density – (area of selected cell x mean fluorescence of background). In addition, to correct for cell autofluorescence, the CTCF values of parasites incubated with L-methionine (negative control) was subtracted from all CTCF values of parasites incubated with AHA.

#### scRNAseq of trypanosomes

After natural infection, skin equivalents were cultured for another 4 h, 12 h, 24 h, and 7 d, respectively. In addition, freshly collected MCFs obtained from tsetse flies were used. Skin equivalents were dissociated and single viable parasites were sorted based on their tdTomato and Calcein AM signal with a FACSAria III cytometer (BD Biosciences) into 48-well plates containing 2.6 µl of 1x lysis buffer (Takara) supplemented with 0.01 µl of RNase inhibitor (40 U/µl; Takara). Immediately after sorting, cells were placed on ice for 5 min and stored at −80 °C.

Library preparation and sequencing was carried out as described previously (Muller et al., 2018). Briefly, 0.2 µl of a 1:20 x 10^6^ dilution of ERCC Spike-in Control Mix 1 (Thermo Fisher Scientific) were added to the lysates of single parasites and libraries were prepared using SMART-Seq v.4 Ultra Low Input RNA Kit (Takara) using a quarter of the reagent volumes recommended by the manufacturer. 27 cycles were used for PCR amplification and cDNA was purified using Agencourt AMPure XP beads (Beckman Coulter) using 15 μl of elution buffer (Takara). Library quantification was performed with a Qubit 3 Fluorometer with dsDNA Hs Assay kit (Life Technologies) and the quality of the libraries was assessed using a 2100 Bioanalyzer with High Sensitivity DNA kit (Agilent). 0.5 ng of cDNA was subjected to a tagmentation-based protocol (Nextera XT, Illumina) using a quarter of the recommended volumes, 10 min for tagmentation at 55 °C and 1 min extension time during PCR amplification. Libraries were pooled and sequencing was performed in paired-end mode for 2 x 75 cycles using Illumina’s NextSeq 500.

#### Analysis of scRNAseq data of trypanosomes

After demultiplexing, data quality was examined using FastQC (version 0.11.7). Illumina adaptors were removed using cutadapt (version 3.2). Trimmed reads were mapped to the TREU927 (version 48) genome with the ERCC spike-in sequences included, using RNASTAR (version 2.6.1b). Read counts for each gene were determined using featureCounts (version 2.0.1) and genes with ≥ 5 aligned reads were considered detected. Only those scRNAseq datasets with > 200 and < 1000 detected genes and a library size between 2 x 10^5^ and 2 x 10^6^ reads were considered for analysis. Principal component analysis (PCA) was performed selecting the 1000 genes with the highest variance. The following libraries were excluded from the analysis due to BARP gene expression: “0h_1_B5_1”, “0h_1_G2”, “0h_1_C5_1”. Differential gene expression analysis was conducted with DESeq2 (version 1.30.0) and SCDE (version 2.18.0; only used in early vs. late analysis). Features with an absolute log2 fold change > 2 and adjusted P value < 0.01 (DEseq2) or a z-score > 1.96 (SCDE) were considered as differentially expressed, respectively.

#### Gene Ontology (GO) Analysis

GO term enrichment within the differential expressed genes between the 5 data sets (MCF, 4 h, 12 h, 24 h, and 7 d) were determined by Fisher’s exact test using the GO enrichment tool on the TriTrypDB webserver (https://tritrypdb.org/tritrypdb/app) (Aslett et al., 2010) and summarized using the REVIGO webtool (Supek et al., 2011) to avoid redundancies. For early vs. late GO term analysis, genes with a z-score > 1.28 (SCDE) were used for enrichment.

#### Tracking of single parasites

Individual trypanosomes were monitored in the skin equivalents based on the signal of the fluorescence reporter tdTomato using a fluorescence stereomicroscope (MZ16 FA, Leica). Movies were acquired for 5 min with a 5x dry objective and a rate of 4 frames per second (FPS). For tracking of single parasites movies were analyzed with Imaris software. Only objects with a diameter larger than 6 µm and slower than 80 µm/s were considered as individual trypanosomes. The maximum gap size was set to 20 frames and tracks shorter than 60 sec were excluded from analysis. All tracks were validated manually and, if necessary, incorrect tracks or incorrectly assigned objects were corrected.

#### Immunofluorescence staining of trypanosomes

Skin equivalents were dissociated and 5 x 10^4^ trypanosomes were sorted on coverslips pre-coated with poly-L-lysine and fixed with 3 % PFA as described above. Parasites were blocked with 500 µl PBS-B for 30 min and incubated with a rat anti-VSG AnTat 1.1 antibody diluted 1:4000 in PBS-B for 1 hour at RT, followed by two wash steps with PBS-B and incubation with a Alexa647-conjugated anti-rat secondary antibody diluted 1:500 in PBS-B for 30 min at RT. Cells were washed twice with PBS-B and coverslips were mounted upside down on a microscope slide with 7.5 µl Fluoromount-G with DAPI. Images were acquired with a DMI6000B wide-field fluorescence microscope (Leica).

#### Scanning electron microscopy

Dermal equivalents or infected skin were fixed in Karnovsky solution (2 % PFA, 2.5 % glutaraldehyde in 0.1 M cacodylate buffer, pH = 7.4) overnight at 4 °C. Specimens were washed 3 times for 5 min at 4 °C with 0.1 M cacodylate buffer, followed by incubation for 1 hour at 4 °C in post-fixation solution (2.5 % glutaraldehyde in 0.1 M cacodylate buffer, pH = 7.4). Samples were washed, incubated in cacodylate buffer containing 2 % tannic acid and 4.2 % sucrose for 1 hour at 4 °C, and washed 3 times with water for 5 min at 4 °C. Subsequently, specimens were dehydrated in an ascending acetone sequence, critical point dried and coated with gold-palladium. Images were acquired with a JEOL JSM-7500F scanning electron microscope using the detector LEI for secondary electrons at 5 kV.

#### Dermal contraction, weight, and viability

To determine dermal contraction and weight loss, the dermal area and weight were measured regularly during culture. Therefore, individual low- and high-density dermal equivalents were cultured in petri dishes (35 mm) containing 3 ml of DMEM supplemented with 10 % FCS, 1 % NEAA, and 1 % penicillin/streptomycin. For up to 3 weeks, each dermal component was photographed once a week using graph paper and the weight was determined with an analytical balance after the medium was completely aspirated. After the measurements, fresh medium was added and the dermal equivalents were cultured further.

Fibroblast viability after compression was determined by fluorescein diacetate (FDA) and propidium iodide (PI) staining. Therefore, dermal equivalents were submersed with 3 ml PBS^-^ containing 20 µM FDA and 3 µM PI for 5 min at 37 °C and subsequently washed 3 times with PBS^-^. Images were acquired using a MZ16 FA fluorescence stereomicroscope (Leica).

#### scRNAseq of skin equivalents

The dermis of two individual hdSEs at day 23 was separated from the epidermis with forceps and subsequently cut into small pieces and incubated in 1 ml of TDB containing 250 µg Liberase TL for 45 min at 37 °C. To isolate NHEK the epidermis was incubated in 3 ml of PBS^-^ with gentle agitation. PBS^-^ was aspirated, the epidermis cut into small pieces and 1 ml of 0.05 % Trypsin/EDTA were added for 15 min at 37 °C. 100 µl of FCS were added and NHEK were released by thorough resuspension. Cells were collected by centrifugation at 300 g for 5 min, resuspended in 500 µl INF medium containing 0.5 µg DAPI and filtered through a 35 µm cell strainer. Viable NHDF and NHEK were sorted based on their negative DAPI signal with a FACSAria III cytometer (BD Biosciences) and collected in a single tube. Cell density was evaluated by microscopy using a hemocytometer and adjusted to 1000 cells/µl.

Subsequently, the Chromium Next GEM Single Cell 3ʹ Reagent Kits v3.1 (10x Genomics) were used for reverse transcription, cDNA amplification, and library construction according to the manufacturers’ instructions. 16.5 µl of cell suspension were used to recover 10,000 cells and libraries were quantified with a Qubit 3 Fluorometer (Thermo Fisher Scientific). Quality was checked using a 2100 Bioanalyzer with High Sensitivity DNA kit (Agilent). Sequencing was performed in paired-end mode with a SP flow cell (2 x 50 cycles) using NovaSeq 6000 sequencer (Illumina).

#### Analysis of scRNAseq data of skin equivalents

CellRanger v3.1.0 (10x Genomics) was used to process scRNAseq data. To generate a digital gene expression (DGE) matrix, reads were mapped to the human reference genome GRCh38 and the number of UMIs for each gene in each cell was recorded.

Seurat R package (version 3.1.4) (Butler et al., 2018) was used for further analysis of scRNAseq data. Only cells with > 2500 genes, between 10^4^ and 10^5^ UMIs, and between 1 % and 15 % mitochondrial (mt) genes were considered for downstream analyses. The “NormalizeData” function was used to normalize sequencing reads of each gene to total UMIs in each cell. The “ScaleData” function was used to scale and center expression levels in the data set for dimensional reduction. The top 2000 features that exhibit high cell-to-cell variation in the dataset were selected with the function “FindVariableFeatures” and used for dimensionality reduction. Total cell clustering was performed by “FindNeighbors” function using the first 20 dimensions and “FindClusters” function at a resolution of 0.4. Dimensionality reduction was performed with “RunTSNE” function and visualized by t-Distributed Stochastic Neighbor Embedding (t-SNE). Marker genes for each cluster were determined with the Wilcoxon rank-sum test by “FindAllMarkers” function. Only those with |‘avg_logFC’| > 0.25 and ‘p_val_adj’ < 0.05 were considered as marker genes.

#### Histological and immunohistochemical staining

Skin equivalents and biopsies of preputial skin were fixed with 4 % PFA at 4 °C overnight, processed for paraffin embedding using a Microm STP 120 tissue processor (Thermo Fisher Scientific), and 5 µm thick sections were prepared using a SM2010 R microtome (Leica). Tissue sections were deparaffinized in xylene and rehydrated using a descending ethanol series. For histological staining, sections were stained with hematoxylin and eosin (H&E) and mounted in Entellan. For immunofluorescence staining, antigens were retrieved by incubation in 10 mM sodium citrate, pH = 6 at 95 °C for 20 min. Sections were permeabilized in PBS^-^ containing 0.3 % Triton X-100 for 5 min and subsequently blocked with PBS-B for 30 min at RT. Next, sections were incubated overnight at 4 °C in a humidity chamber with the appropriate primary antibody, followed by washing 3 times for 5 min with PBS^-^ containing 0.5 % Tween-20 (PBS-T). Tissue sections were incubated with the appropriate secondary antibody for 1 hour at RT in the dark. After three washing steps with PBS-T for 5 min, sections were mounted in Fluoromount-G with DAPI. All images were acquired with a BZ-9000 fluorescence microscope (Keyence). The thickness of the cellular epidermis was measured with ImageJ/Fiji in H&E-stained sections of skin equivalents.

#### Rheology

Mechanical properties of high- and low-density dermal equivalents were analyzed with a MCR 102 rheometer (Anton Paar). Pieces with a diameter of 8 mm were punched out of individual dermal equivalents using an 8 mm tissue punch. Measurements were performed using a plate-plate geometry of 8 mm diameter and a gap size of 1 mm at RT. Amplitude sweeps from 0.01 % to 100 % strain deformation at 1 Hertz were performed to investigate viscoelastic properties.

### Quantification and Statistical Analysis

The number of individual experiments is annotated in corresponding figure legends. Statistical significance was determined using the unpaired Student’s t-test or Mann-Whitney U-test in GraphPad Prism version 7 (GraphPad). P values correlate with symbols as follows: ns = not significant, p > 0.05, ∗ p ≤ 0.05, ∗∗ p ≤ 0.01, ∗∗∗ p ≤ 0.001, ∗∗∗∗ p ≤ 0.0001.

### Data and Software Availability

The accession number for single cell trypanosome RNA sequencing data is NCBI GEO: GSE174198 The accession number for single cell high-density skin equivalent RNA sequencing data is NCBI GEO:

## Supplemental item titles and legends

**Table S1. Related to Figure 2.** scRNAseq analysis of skin equivalents.

**Table S2. Related to Figure 5.** scRNAseq analysis of trypanosomes.

**Table S3. Related to Figure 7.** scRNAseq analysis of trypanosomes.

**Table S4. Related to Figure 5 and 7.** GO term enrichment of trypanosomes.

**Movie 1. Related to Figure 3.** Natural transmission of *T. brucei* parasites to skin equivalents by tsetse fly. Three hdSEs were stacked on top of each other and subsequently tsetse flies with a mature salivary gland infection were allowed to bite into the stack.

**Movie 2: Related to Figure 4.** Tracking of trypanosomes in skin equivalents at various timepoints post-infection. Skin equivalents were infected with *T. brucei* parasites by tsetse flies and cultured for 3 days. At the times indicated individual trypanosomes were recorded using a fluorescence stereomicroscope. Movies were acquired for 5 min at 4 fps and analyzed with Imaris software. Video plays at 20x speed.

**Movie 3. Related to Figure 6.** Skin-resident trypanosome at 7 dpi expressing the stumpy marker PAD1. Skin equivalents were infected with *T. brucei* parasites by tsetse flies and cultured for 7 days. The movie was recorded with a fluorescence stereomicroscope for 1 min at 4 fps. Video plays at 3x speed.

## Supplementary figures

**Figure S1. Related to Figure 1.**
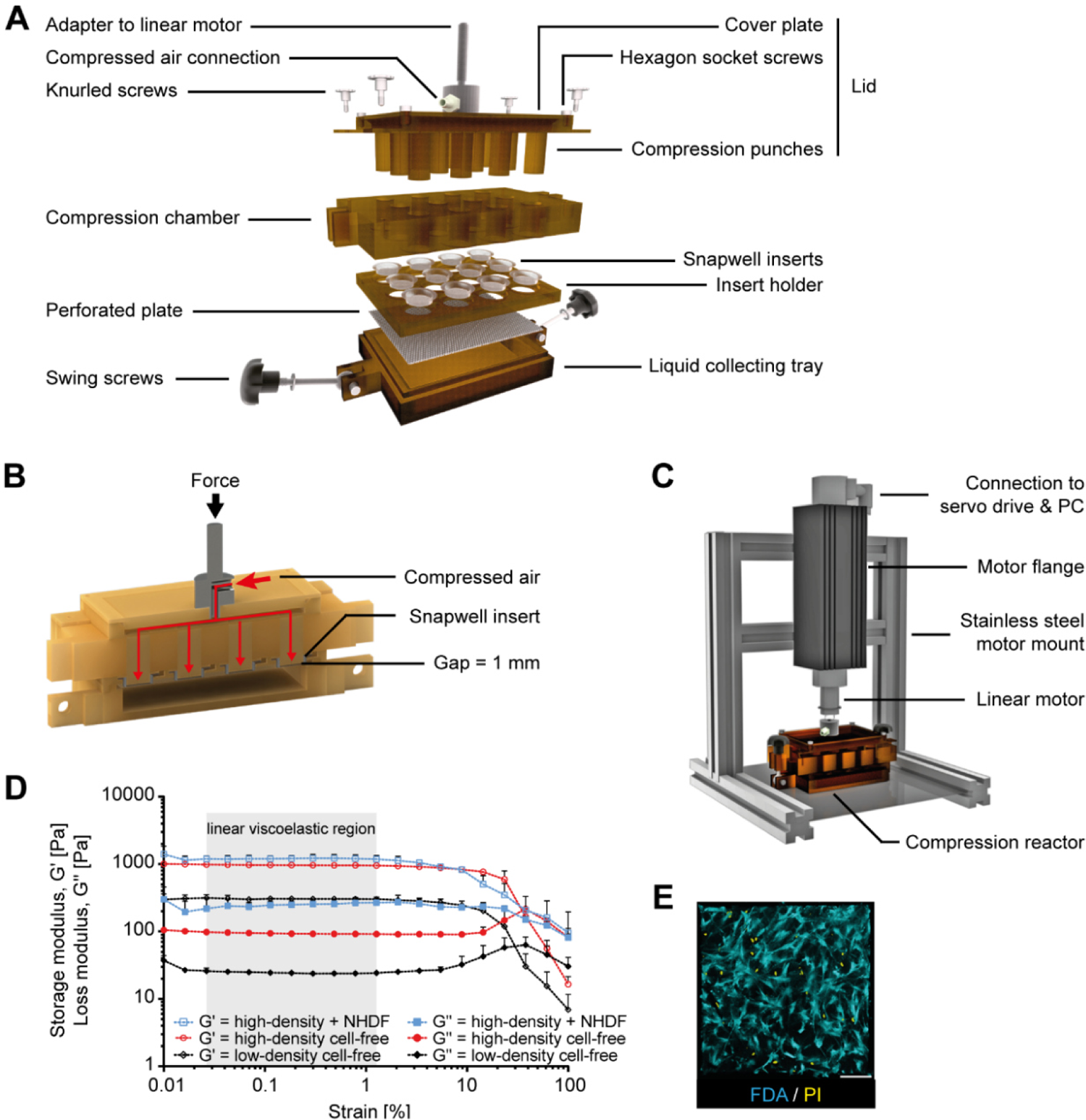
Bioreactor design and compression system. **(A)** Representation of the computer-aided design (CAD) of the compression reactor with all individual parts. **(B)** Cross section through the closed reactor after complete compression. The red arrows mark the route of the compressed air through the reactor. The compressed air was introduced through the compressed air connection, which could then exit through the channels in the individual compression punches to release the dermal equivalents. **(C)** Representation of the compression system. Shown is the configuration of the linear motor, which was attached to a stainless steel motor mount via the motor flange. The motor can be controlled with the LinMot®-Talk software by connecting it to a computer. **(D)** Plot of oscillatory rheological measurements. Storage, G′ (open circles/squares), and loss modulus, G′′ (filled circles/squares), of cell-free and NHDF-populated high- and low-density dermal equivalents was determined in dependency on the strain deformation (ω = 6.28 rad/s, T = 22 °C). The linear viscoelastic region (gray box) was determined according to DIN 53019-4. Data represent mean ± SD (n = 3 - 6). **(E)** Assessment of NHDF viability in high-density dermal equivalents on day 23 after compression. NHDF were stained with fluorescein diacetate (FDA, cyan) and propidium iodide (PI, yellow) for live/dead discrimination. Scale bar, 100 µm.

**Figure S2. Related to Figure 2.**
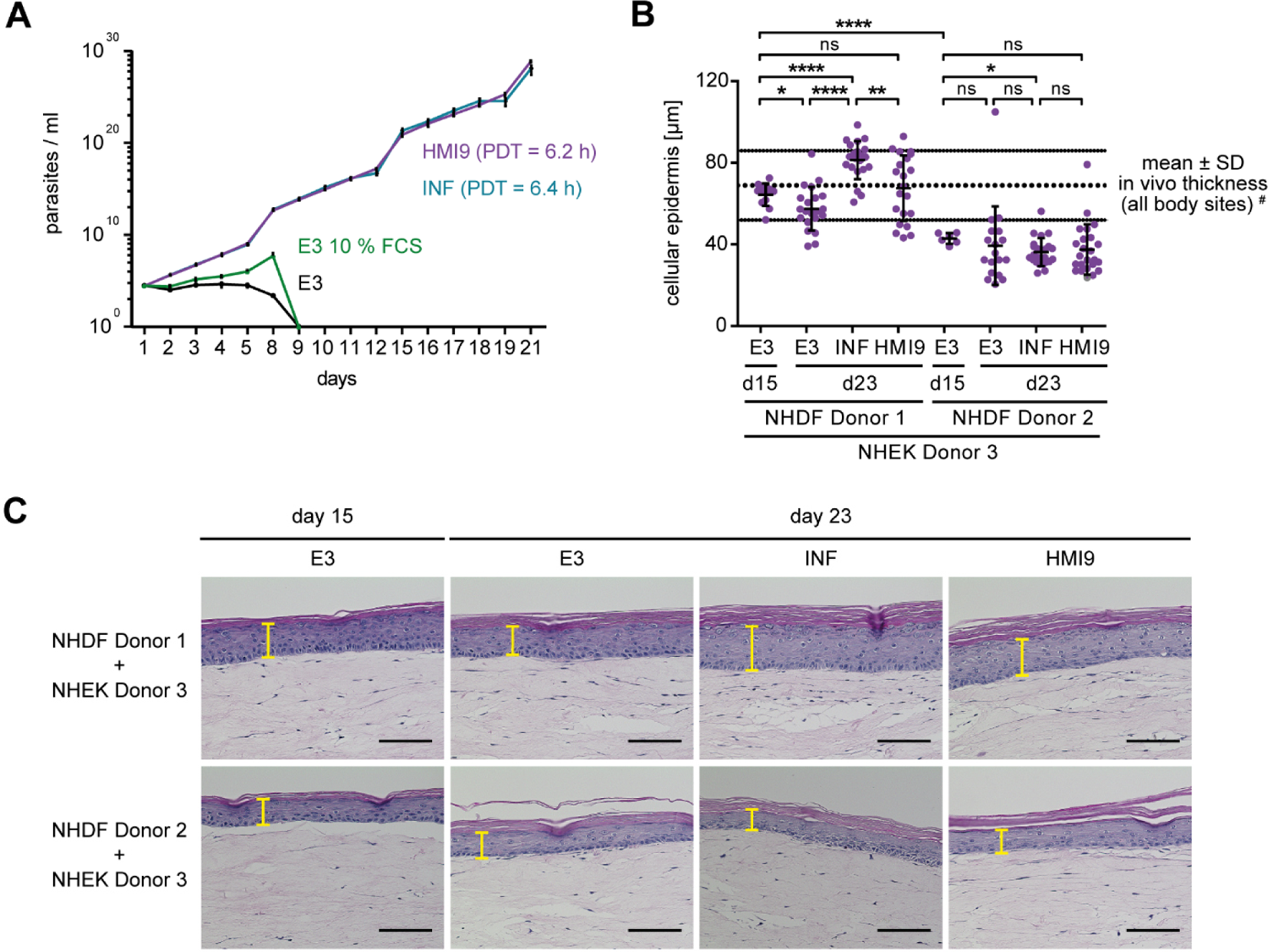
*T. brucei* parasites and high-density skin equivalents can be cultured with the same medium without growth deficits. **(A)** Growth and population doubling times (PDT) of monomorphic Lister 427 90:13 parasites in various media: trypanosome medium HMI9 (purple), skin medium E3 supplemented with 10 % FCS (green) or without FCS (black), and a mixture of both media (= INF, cyan). Data represent mean ± SD (n = 3). **(B)** Analysis of the thickness of the cellular epidermis of hdSEs generated with NHDFs from two different donors (donor 1 and 2) and NHEKs from a third donor (donor 3). Skin equivalents were cultured with E3 medium until day 15. On day 15, the medium was changed to E3, INF, or HMI9 until day 23. ImageJ was used to measure the thickness of the cellular epidermis in hematoxylin and eosin-stained cross sections of hdSEs. Data represent mean ± SD. Unpaired t-test, **** p < 0.0001, ** p < 0.01, * p < 0.05, ns, not significant. # (Sandby-Moller et al., 2003). **(C)** Hematoxylin and eosin-stained cross sections of hdSEs cultured with E3, INF, or HMI9 media at day 23 compared with day 15. The yellow markings indicate the thickness of the cellular epidermis. Scale bar, 100 µm.

**Figure S3. Related to Figure 2.**
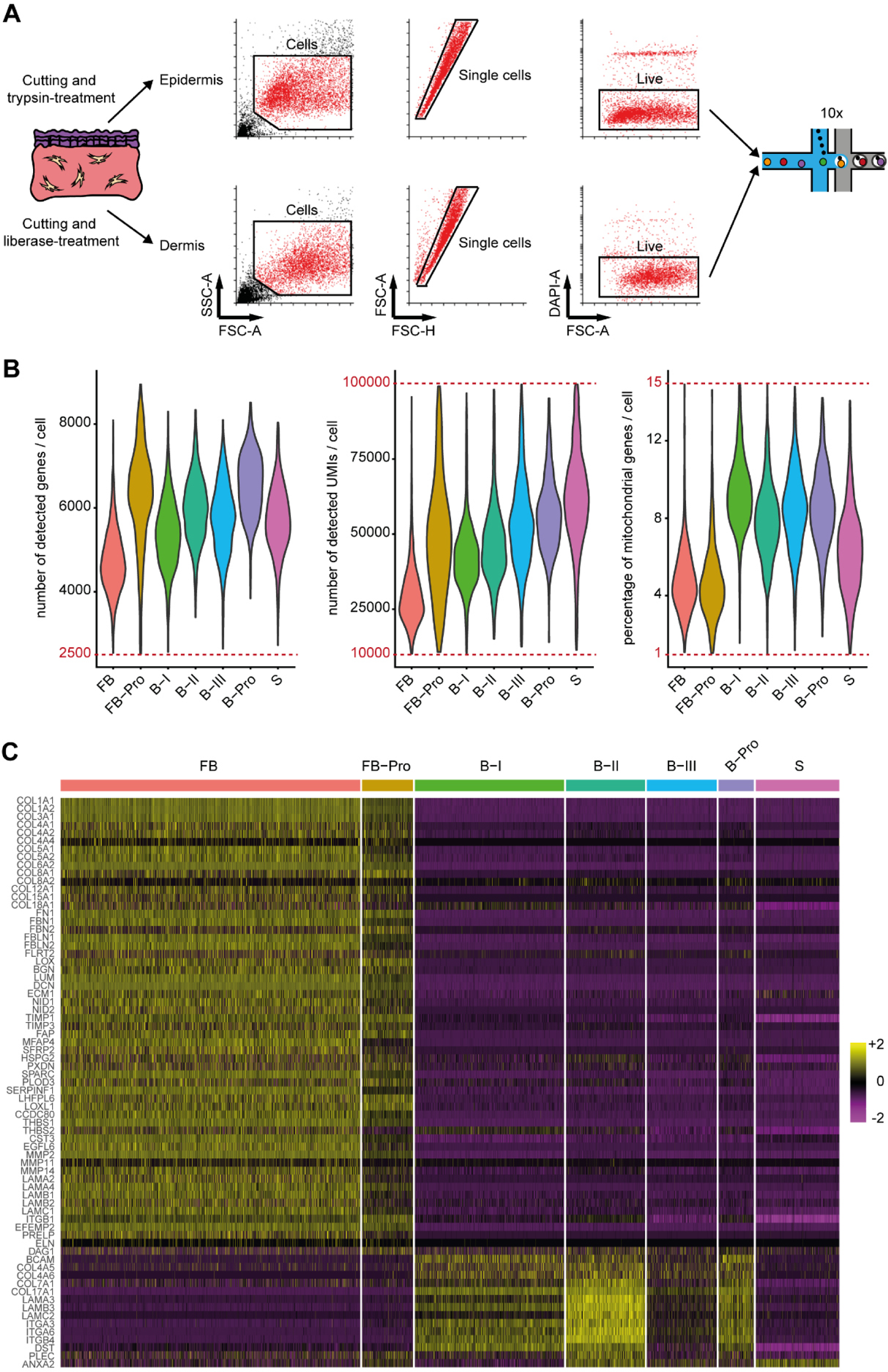
Strategy and quality control of the single-cell RNA sequencing experiment of high-density skin equivalents and analysis of the expression of extracellular matrix-associated genes. **(A)** The epidermis and dermis of hdSEs were separated, cut, and dissociated by trypsin or liberase treatment. Cell suspensions were separated by FACS and single, live cells were sorted. Subsequently, cells were loaded on the droplet-based 10x platform according to the manufacturers’ instructions and sequenced. **(B)** Quality control of 10x data showing the number of detected genes, UMIs (unique molecular identifier), and the percentage of mitochondrial (mt) genes per cell in each of the 7 subclusters. The red lines in each graph represent quality control thresholds and cells above or below these thresholds were excluded from analysis. **(C)** Heatmap showing the scaled expression levels of extracellular matrix-associated genes differentially expressed in each cluster. The color key from pink to yellow indicates low to high gene expression levels. Each column represents a single cell, each row represents an individual gene. Gene names are listed to the left.

**Figure S4. Related to Figures 3 and 4.**
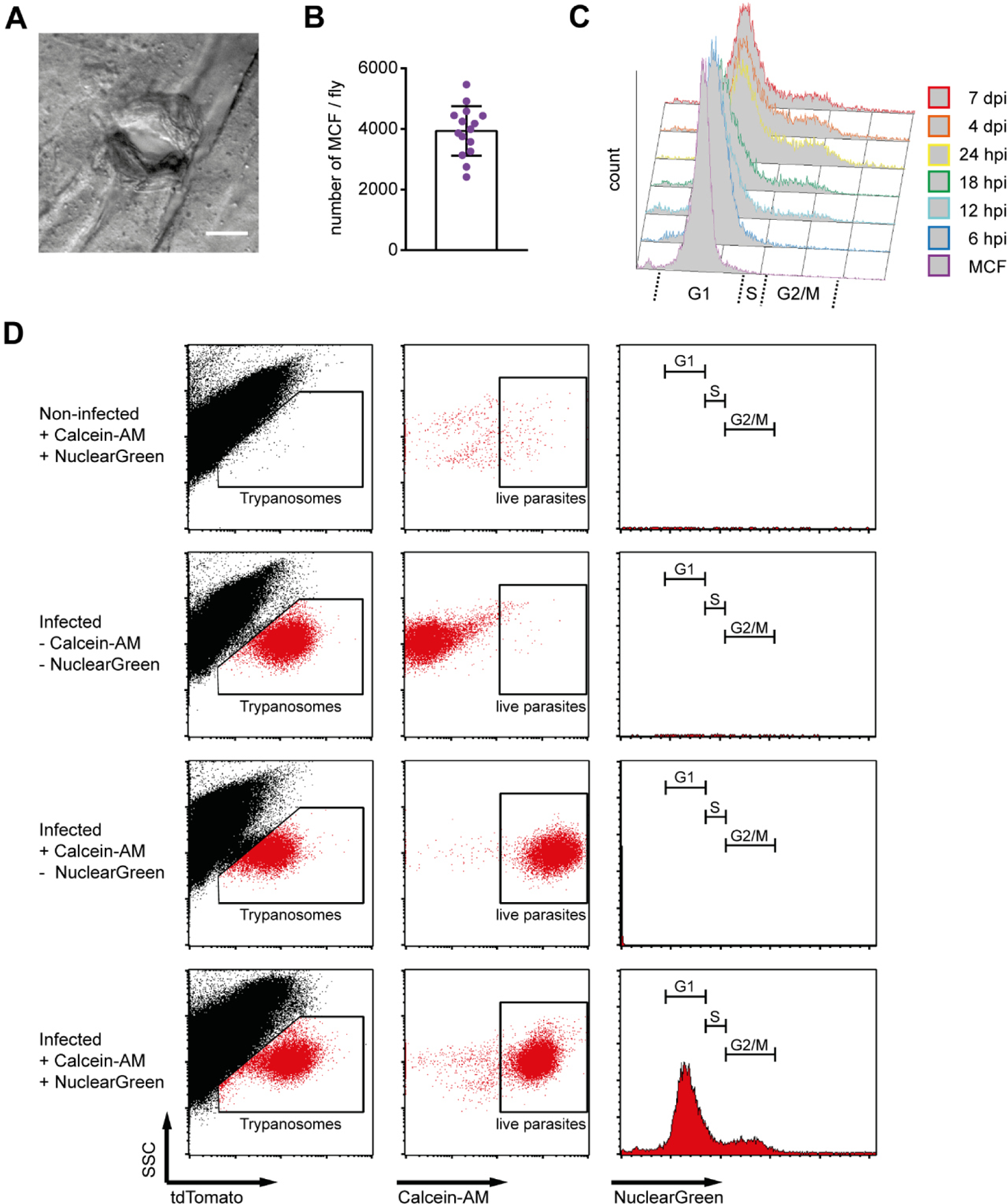
Epidermal wound, number of tsetse-transmitted parasites and FACS gating strategy. **(A)** Stereo microscopy of the epidermal wound caused by the fly bite. Scale bar, 50 µm. **(B)** Number of metacyclic forms (MCF) in spit samples from individual salivary gland-positive flies. The maximum number of injectable MCFs per fly bite was 3934 on average. Results are shown as mean ± SD (n = 15). **(C)** Representation of cell cycle histograms of the flow cytometric assessment of the cell cycle distribution of skin-residing trypanosomes at various timepoints post-infection. **(D)** Flow cytometric profiles showing the gating strategy to detect trypanosomes (tdTomato positive) isolated from skin equivalents, to determine parasite viability (Calcein-AM positive), and to assess cell cycle distribution with NuclearGreen staining.

**Figure S5. Related to Figure 5.**
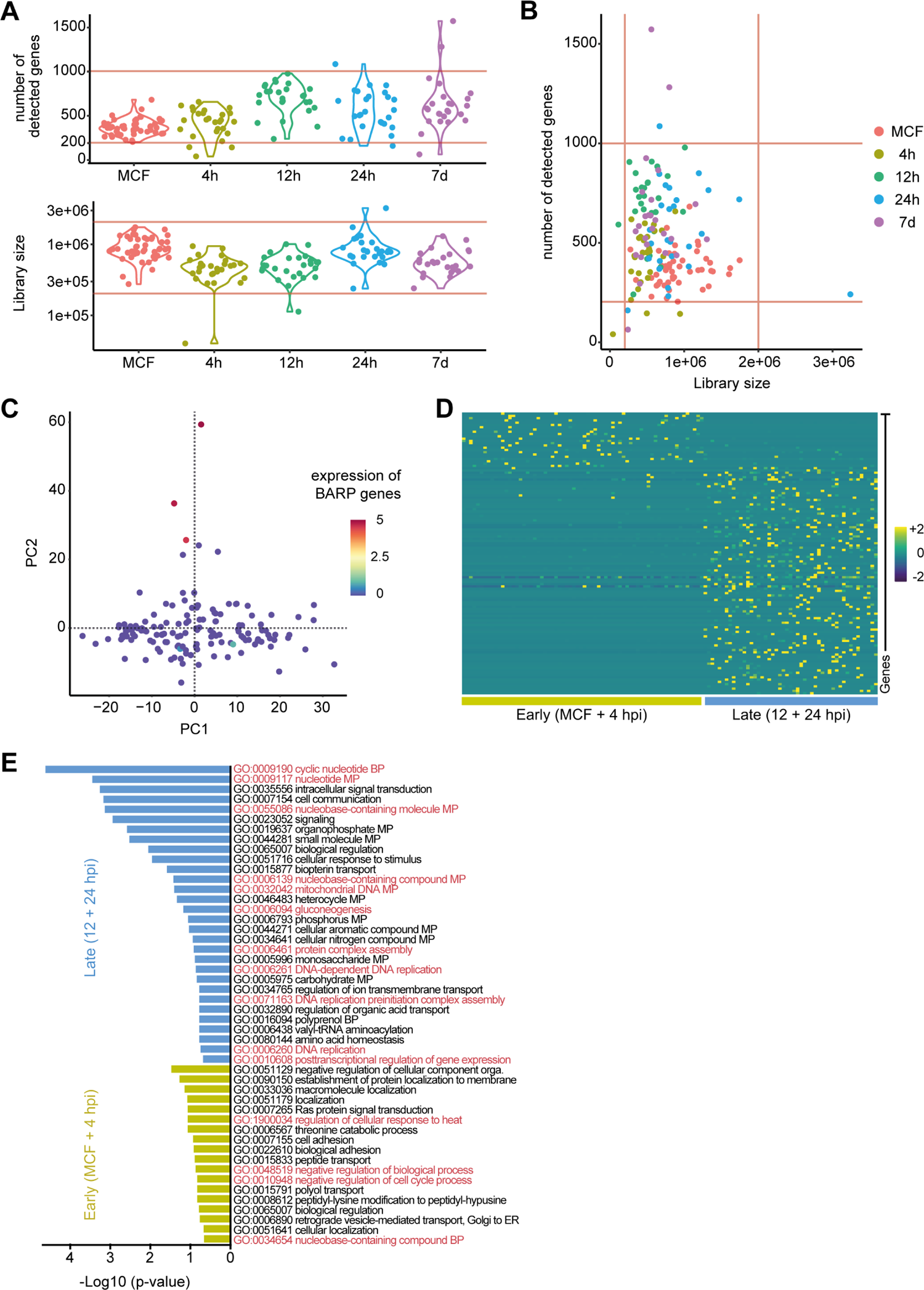
Quality control of the single-parasite RNA sequencing experiment. **(A, B)** Violin and dot plots showing the number of genes detected across all libraries and library size of individual *T. brucei* parasites (MCF, n = 48; 4 h, n = 27; 12 h, n = 27; 24 h, n = 26; 7 d, n = 25). The red lines represent the thresholds used for quality control. **(C)** Feature plot of the 142 transcriptomes that passed quality control showing the normalized expression of fly-stage specific BARP genes (“Tb927.9.15510”, “Tb927.9.15520”, “Tb927.9.15530”, “Tb927.9.15540”, “Tb927.9.15550”, “Tb927.9.15560”, “Tb927.9.15570”, “Tb927.9.15580”, “Tb927.9.15590”, “Tb927.9.15600”, “Tb927.9.15610”, “Tb927.9.15620”, “Tb927.9.15630”, “Tb927.9.15640”). The color key from blue to red indicates low to high gene expression levels. **(D)** Heatmap showing the scaled expression levels of upregulated genes of early (MCFs + 4 hpi) and late (12 hpi + 24 hpi) timepoints, as identified by SCDE (log2 fold change > 2, z-score > 1.96). Each column represents a single parasite, each row represents an individual gene. The color key from purple to cyan indicates low to high gene expression levels. **(E)** Biological process-associated GO terms significantly enriched in early (MCFs + 4 hpi) and late (12 hpi + 24 hpi) timepoints. GO terms were filtered with the Revigo webtool to avoid redundancies. Bars represent the level of significance of the term enrichment.

**Figure S6. Related to Figure 7.**
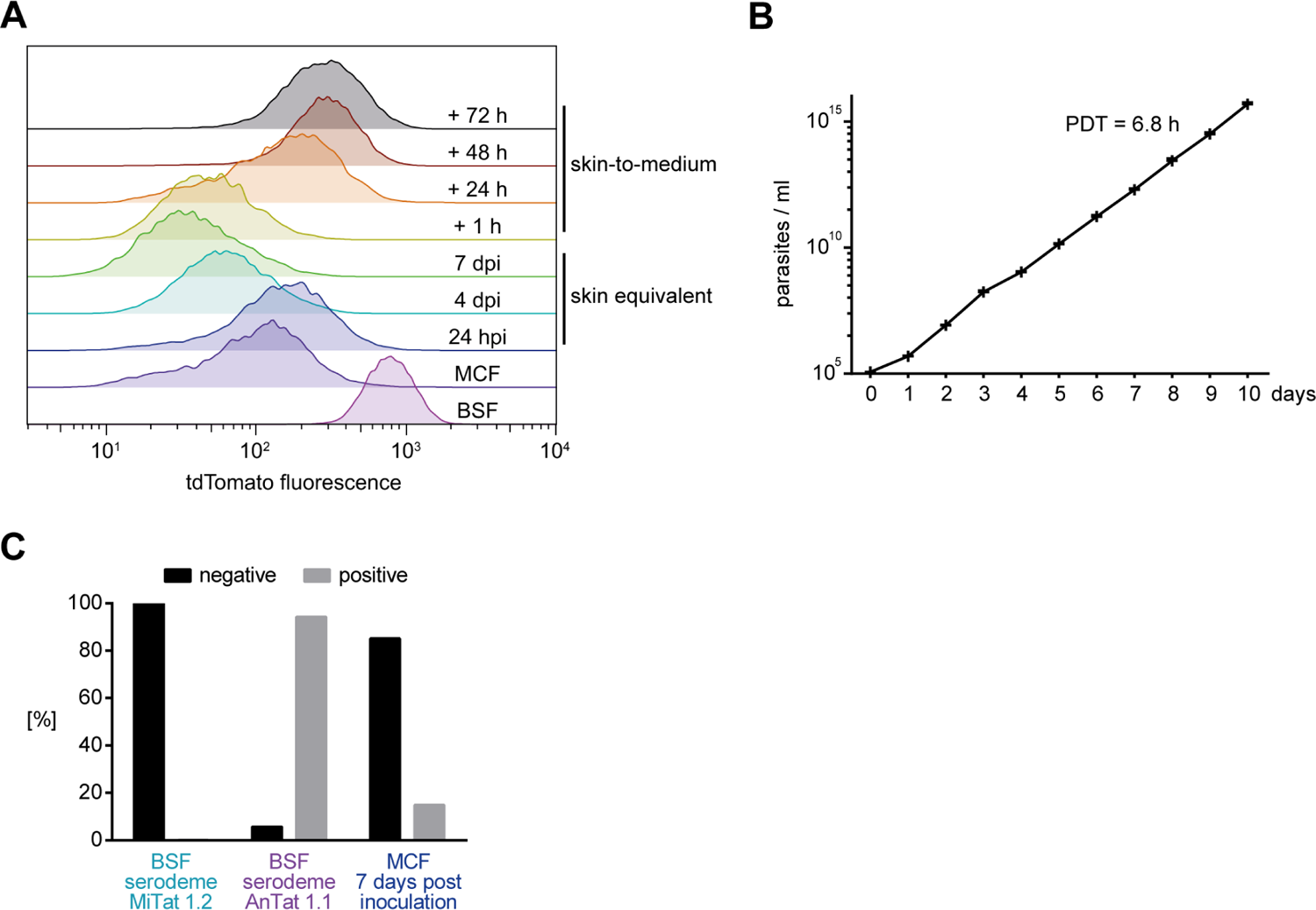
Mean fluorescent intensity of tdTomato, and growth and VSG AnTat 1.1 expression of metacyclic trypanosomes under axenic conditions. **(A)** Representation of flow cytometric histograms of the tdTomato fluorescence intensities of skin-residing trypanosomes compared to BSFs, MCFs, and skin-parasites at 7 dpi, which were transferred to culture medium for up to 3 days (skin-to-medium). **(B)** Growth and population doubling time (PDT) of metacyclic trypanosomes inoculated into HMI9 medium supplemented with methylcellulose. MCFs were harvested from tsetse flies with a mature salivary gland infection by salivation. Data represent mean ± SD (n = 3). **(C)** Immunocytological quantification of VSG AnTat 1.1 expression of monomorphic BSFs of the serodeme MiTat 1.2 compared with pleomorphic BSFs of the serodeme AnTat 1.1 and MCFs inoculated for 7 days into HMI9 medium supplemented with methylcellulose. MCFs were obtained from tsetse flies infected with stumpy forms of the serodeme AnTat 1.1. The percentage of parasites which stained negative or positive for AnTat 1.1 are shown. 30,000 parasites were analyzed per condition in a single experiment.

